# Amoeboid-like neuronal migration ensures correct horizontal cell layer formation in the developing vertebrate retina

**DOI:** 10.1101/2021.10.15.464510

**Authors:** Rana Amini, Raimund Schlüßler, Stephanie Möllmert, Archit Bhatnagar, Jochen Guck, Caren Norden

**Affiliations:** Instituto Gulbenkian de Ciência, Rua da Quinta Grande 6, 2780-156 Oeiras, Portugal; Max Planck Institute of Molecular Cell Biology and Genetics, Pfotenhauerstraße 108, 01307 Dresden, Germany; Biotechnology Center, Center for Molecular and Cellular Bioengineering, Technische Universität Dresden, 01307 Dresden, Germany; Max Planck Institute for the Science of Light and Max-Planck-Zentrum für Physik und Medizin, 91058 Erlangen, Germany; Physics of Life,Technische Universität Dresden, 01307 Dresden, Germany

**Keywords:** Retina, Neuronal Migration, Horizontal Cells, Amoeboid Cell Migration, Neuronal lamination, Zebrafish

## Abstract

As neurons are often born at positions different than where they ultimately function, neuronal migration is key to ensure successful nervous system development. Radial migration during which neurons featuring unipolar and bipolar morphology, employ pre-existing processes or underlying cells for directional guidance, is the most well explored neuronal migration mode. However, how neurons that display multipolar morphology, without such processes, move through highly crowded tissue environments towards their final positions remains elusive. To understand this, we here investigated multipolar migration of horizontal cells in the zebrafish retina. We found that horizontal cells tailor their movements to the environmental spatial constraints of the crowded retina, by featuring several characteristics of amoeboid migration. These include cell and nucleus shape changes, and persistent rearward polarization of stable F-actin, which enable horizontal cells to successfully move through the crowded retina. Interference with the organization of the developing retina by changing nuclear properties or overall tissue architecture, hampers efficient horizontal cell migration and layer formation. Thus, cell-tissue interplay is crucial for efficient migration of horizontal cells in the retina. In view of high proportion of multipolar neurons, the here uncovered ameboid-like neuronal migration mode might also be crucial in other areas of the developing brain.

## INTRODUCTION

Migration of newly-born neurons to their designated positions is a key step in the establishment of neural circuits and thereby function of the central nervous system (CNS). A variety of migration modes have been uncovered featuring diverse cell morphologies ranging from unipolar or bipolar to multipolar (Marin and Rubenstein 2003, Tabata and Nakajima 2003, Kawaji et al. 2004, Ayala, Shu and Tsai 2007, Tanaka et al. 2009, Cooper 2013, Rahimi-Balaei et al. 2018, Gressens 2000). As of yet, most research unveiling the cellular and molecular mechanisms of neuronal migration has focused on radial migration which is limited to movements of unipolar and bipolar neurons, perpendicular to the tissue surface (Angevine and Sidman 1961, Berry and Rogers 1965, Morest 1970b, Morest 1970a, Rakic 1971, Walsh and Cepko 1988, Marin and Rubenstein 2001, Nadarajah et al. 2001, Nadarajah and Parnavelas 2002, Cooper 2013).

During radial migration, migrating neurons determine their direction of movement by two different strategies: 1) sending process(es) which anchor the migrating neurons to the tissue lamina(e), and thereby facilitating the faithful arrival of neurons at their destination (somal translocation) (Nadarajah et al. 2003), or 2) moving along radially-oriented fibers of neural progenitors known as radial glia, that provide physical scaffolding for migrating neurons (glia-guided migration) (Rakic 1971, Rakic 1972, Edmondson and Hatten 1987, Hatten 1990, Gertz and Kriegstein 2015). In both scenarios, the migrating neurons exhibit elongated unipolar or bipolar morphologies in the direction of travel and move unidirectionally via radial paths to their final positions (Nadarajah et al. 2001, Nadarajah et al. 2003).

In contrast, neurons that display multipolar morphology are neither attached to the tissue lamina(e) nor move along radial glia fibers and instead extend multiple dynamic processes in various directions (Stensaas 1967, Shoukimas and Hinds 1978, Nowakowski and Rakic 1979, Gadisseux et al. 1990, Tabata and Nakajima 2003, Honda, Tabata and Nakajima 2003, Noctor et al. 2004, Cooper 2014). Examples include interneurons of the mammalian neocortex (Nadarajah et al. 2003, Tabata and Nakajima 2003, Tanaka et al. 2006, Tanaka et al. 2009), pyramidal and granule neurons in the mammalian hippocampus (Kitazawa et al. 2014, Namba, Shinohara and Seki 2019). How multipolar neurons, without any predisposed migratory information or scaffolds reach their destination during brain development is much less explored. It is particularly not known whether and how multipolar neurons adjust their migration mode, path and efficiency to their densely-packed surrounding environment (Bondareff and Narotzky 1972, Sekine et al. 2011). This knowledge gap is partly due to the inaccessibility of many brain regions for *in vivo* imaging.

One CNS region that allows for such imaging approaches in a quantitative manner is the zebrafish retina (Galli-Resta et al. 2008). The retina consists of five major neuron types; photoreceptor (PR), horizontal cell (HC), bipolar cell (BC), amacrine cell (AC), retinal ganglion cell (RGC), and a single glial cell-type; Müller Glia (MG). During development, retinal neurons move from their birth-site to their functional residence **(Fig 1A)**, and reproducibly assemble into three distinct nuclear layers; outer nuclear layer (ONL), inner nuclear layer (INL), ganglion cell layer (GCL). Synapses between these nuclear layers form at two nuclei-free layers, known as plexiform layers; the outer plexiform layer (OPL) and the inner plexiform layer (IPL) **(Fig 1A’)**.

**FIGURE 1:**
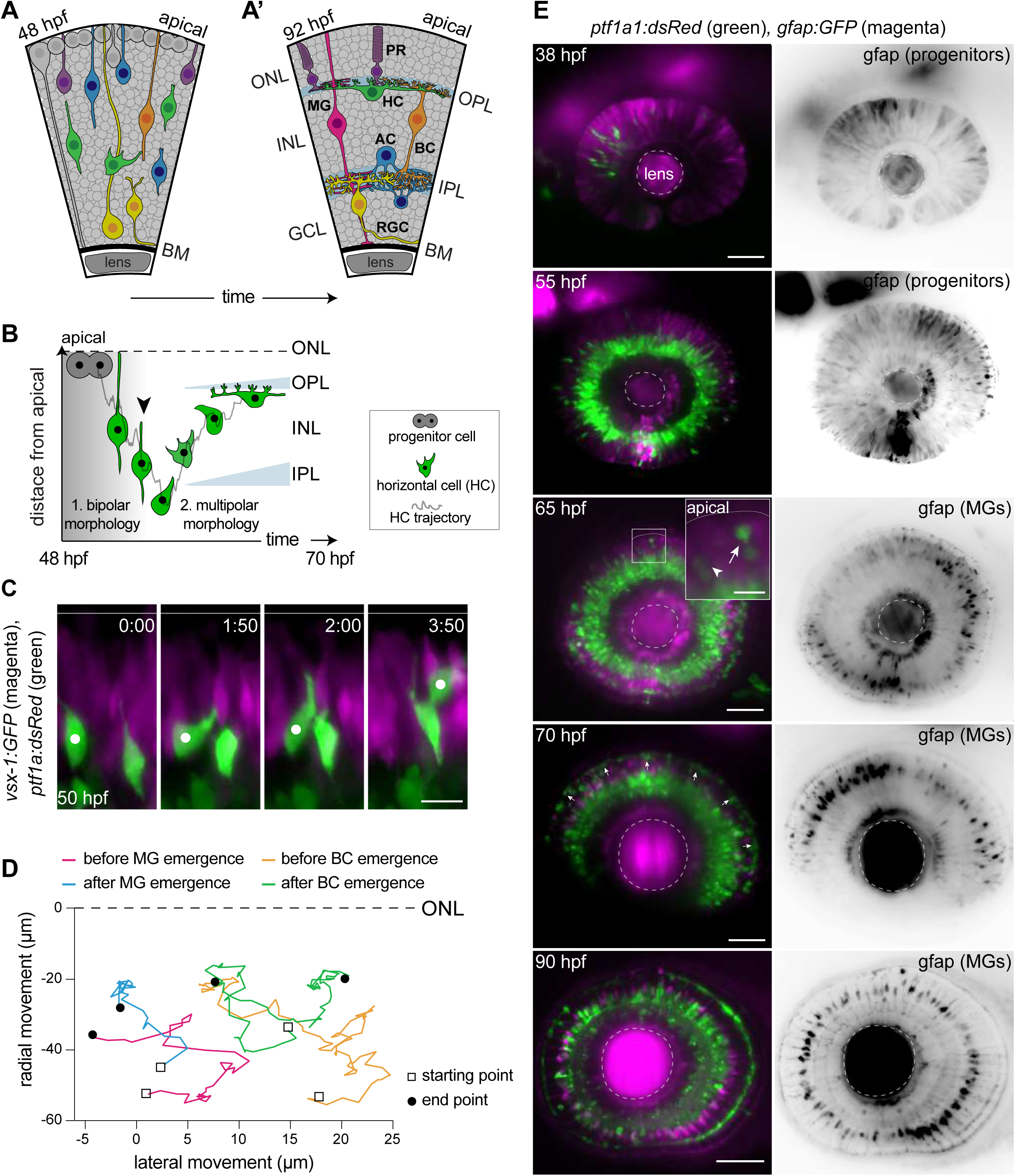
Migrating HCs do not use radially-oriented MGs or BCs for guidance. **(A-A’)** Schematic representation of zebrafish retinal development: **(A)** Birth and neuronal migration (48 hpf). Neuroepithelial progenitors [gray] divide in the apical side and give birth to five major neuron types; photoreceptor (PR), horizontal cell (HC), bipolar cell (BC), amacrine cell (AC), retinal ganglion cell (RGC). Different neuron types are on their way to their appropriate positions. **(A’)** The layered organization of the 92 hpf zebrafish retina. Retinal neurons and a single type of retina glial cell; Müller Glia (MG) are arranged in three nuclear-layers from apical to basal: outer nuclear layer (ONL), inner nuclear layer (INL), ganglion cell layer (GCL). These layers are separated by two synaptic layers: outer plexiform layer (OPL) and inner plexiform layer (IPL). The basement membrane (BM) separates the GCL from the vitreous body. **(B)** Scheme of bidirectional and bimodal HC migration. Arrowhead: detachment from the apical surface and onset of multipolar migration. Gray line: a typical HC migration trajectory (stills and trajectory in Supp Fig 1A-B). **(C)** Time-lapse of HC tangential migration before BC lamination (50 hpf). White dot: tracked HC. *Tg(vsx-1:GFP)* labels BCs [magenta] and *Tg(ptf1a:DsRed)* marks HCs [green]. Time in h:min. Scale bar: 10 μm **(D)** Basal-to-apical migration trajectories of HCs before and after emergence of mature radially-oriented BCs and MGs. HCs use both radial (apical-basal) and tangential (lateral) routes to move within INL. **(E)** Maximum projections of immunofluorescence retinae at different developmental stages: 38, 55, 65, 70 and 90 hpf. *Tg(ptf1a:dsRed)* is expressed in HCs and ACs [green] and *Tg(gfap:GFP)* labels MGs [magenta]. **38 hpf:** birth of and initiation of HC basal migration. GFAP^+^ cells are progenitors. **55 hpf:** initiation of HC apical migration. **65 hpf:** peak of HC apical migration. Higher magnification inset of the outlined region shows an HC at its final destination [arrow], and a migrating HC most likely *en route* to the apical [arrowhead]. **70 hpf:** emergence of HC layer and MG maturation. Arrowheads show HCs within the HC layer. Scale bars: 50 μm, inset 65 hpf: 10 μm.

Retinal neurons follow diverse and complex migratory modes and routes to reach their destinations (Chow et al. 2015, Icha et al. 2016a, Amini, Rocha-Martins and Norden 2017, Amini, Labudina and Norden 2019, Rocha-Martins et al. 2021). So far however, parameters that influence successful migration of the diverse neuron types in their complex and highly dynamic environment are only beginning to be understood (Chow et al. 2015, Icha et al. 2016a, Amini et al. 2019, Rocha-Martins et al. 2021).

Especially intriguing is the movement of HCs, the retinal interneurons that modulate information flow from PRs to BCs (Chaya et al. 2017). HCs exhibit a bidirectional and bimodal migration pattern that features a switch from bipolar to multipolar morphology (Edqvist and Hallbook 2004, Weber et al. 2014, Chow et al. 2015, Amini et al. 2019) **(Fig 1B, Sup-Fig 1A, Video 1)**. Newborn HCs display a bipolar morphology and migrate radially from their apical birth-site to the center of the INL while keeping an attachment to the apical surface (Phase 1). Upon detachment of this anchorage (**Sup-Fig 1C, Video 2**), HCs acquire a multipolar morphology and migrate with frequent direction changes (Phase 2) (Chow et al. 2015, Amini et al. 2019). During this phase, HCs move deeper into the INL before turning apically towards the HC layer at the upper border of the INL, beneath the OPL **(Fig 1B, Sup-Fig 1A-B)**.

The developing zebrafish retina is a densely-packed environment (Matejcic, Salbreux and Norden 2018) which undergoes structural changes in space and time during neuronal lamination. How multipolar HCs adapt their migration behavior and trajectories to the changing and crowded microenvironment of the retina, remains unexplored. The fact that HCs follow unpredictable migration paths despite an overall directionality (Amini et al. 2019) implies that HC path selection is not intrinsically programmed but that the surrounding environment plays a key role. However, if, how, and to what extend cellular and tissue-wide properties influence HC movements towards layer formation remained unexplored.

We here investigated the cellular and tissue-wide parameters that influence HC migration in the developing zebrafish retina. We show that HCs constantly tailor their migration behavior to the limited space within the densely-packed retina by frequent and reversible amoeboid-like shape changes. We further uncover that changing organization of the developing retina at the single-cell or tissue-scale impairs efficient HC migration and perturbs proper HC layer formation.

## RESULTS

### HORIZONTAL CELLS DO NOT EMPLOY GLIA-GUIDED MIGRATION

It was shown that HCs lose their apical attachment **(Sup-Fig 1C, Video 2)** ∼2-3 hrs after apical birth and subsequently display multipolar morphology when moving towards their final destination (Chow et al. 2015, Amini et al. 2019). Since many multipolar interneurons in the neocortex use radially-oriented cells featuring bipolar morphology as their migratory scaffold (Cooper 2014), we asked if such a phenomenon is also seen for HCs in the retina. To this end, we examined whether migrating HCs move along BCs or MGs, the radially-oriented retinal cells with bipolar morphology **(Fig 1A’).**

We performed light-sheet time-lapse imaging using double-transgenic zebrafish embryos *Tg(vsx1:GFP) x Tg(Ptf1-a:dsRed)* labeling BCs and HCs, respectively. The fact that neuronal differentiation in the retina occurs in a wave-like manner (Hu and Easter 1999), allowed us to simultaneously visualize retinal regions hosting immature (unlaminated) and mature (laminated) BCs **(Sup-Fig 1D)**. We noted that in regions without laminated BCs, HCs were either already at **(Sup-Fig 1D)** or en route towards their final position. Further, many HCs did not follow strictly radial migratory trajectories but instead moved in all three dimensions while frequently changing direction **(Fig 1C)**. The same was seen in regions that hosted laminated BCs **(Fig 1D, Video 3)**, arguing against steady, direct interaction between radially-oriented BCs and migrating HCs. Similar observations were made when probing a possible association between migrating HCs and developing MGs, the glial cells of the retina. Immunofluorescence stainings of double-transgenic animals *Tg(gfap:GFP)* x *Tg(Ptf1-a:dsRed)* marking MGs and HCs, showed that prior to MG emergence at 48 hpf, GFAP (glial fibrillary acidic protein) was expressed in radially-oriented retinal neurogenic progenitors (Bernardos and Raymond 2006, Rapaport et al. 2004) **(Fig 1E, 38 hpf)**. No GFAP^+^ cells featuring mature MG morphology at the onset (48 hpf) or peak (55-65 hpf) of HC migration were observed **(Sup-Fig 1D, 50 hpf)**. When mature MGs emerged (around 65 hpf), some HCs were still *en rout*e towards the apical side **(Figure 1E, 65 hpf)**, while others had already reached the prospective HC layer **(Figure 1E, 65 hpf)**. From 70-72 hpf, when GFAP was specifically expressed in mature MGs (MacDonald et al. 2015) **(Fig 1E, Sup-Fig 1E)**, the majority of HCs were already integrated into the HC lamina **(Figure 1E, 70 hpf)**. As seen in embryos expressing BC markers, also in embryos labelled for MGs, HCs followed tangential routes perpendicular to the radial orientation of MG fibers, both before and after MG maturation **(Fig 1D, Sup-Fig 1E’-E”)**. Thus, we conclude that HCs are unlikely to move along the radially-oriented BC or MG fibers.

### INNER NUCLEAR LAYER IS A DENSELY-PACKED TISSUE ENVIRONMENT NOT DOMINATED BY ECM

Cells migrating within tissues can be influenced by mechanical cues from their environment. For example, local gradients in extracellular matrix (ECM) stiffness can guide cell migration in a process termed durotaxis (Lo et al. 2000, Isenberg et al. 2009, Roca-Cusachs, Sunyer and Trepat 2013, Bollmann et al. 2015). In the context of neuronal migration, the ECM can act either as an instructive scaffold along which migration occurs or as a barrier for migrating neurons (Franco and Muller 2011). We thus asked whether ECM components could influence HC migration, focusing on Laminin α1, a glycoprotein that forms fibrous structures (Timpl et al. 1979, Chung et al. 1979), and Neurocan (termed here ssNcan in *Tg(ubi: ssNcan-EGFP)*) as a biosensor for Hyaluronic acid (HA) (De Angelis et al. 2017, Grassini et al. 2018) a glycosaminoglycan forming hydrogel-like structures. Our immunofluorescence imaging showed that anti-Laminin α1 and HA were only detected in the basement membrane of the retina at all developmental stages **(Fig 2A-B, arrowheads)** and never within the INL wherein HC migration takes place **(Fig 2A-B)**. Thus, it is unlikely that HCs use ECM scaffolds as a main migratory substrate to reach their destination. We next asked whether the developing zebrafish retina features mechanical gradients during HC migration, and if yes, whether and how these change in space and time. To address this point, we used Brillouin light scattering microscopy, a non-invasive technique which has been recently applied to a broad range of living biological systems including different areas of the CNS (Girard et al. 2015, Schlussler et al. 2018, Scarcelli and Yun 2012, Scarcelli, Kim and Yun 2011, Mattana et al. 2017). This technique provides information about tissue compressibility by measuring the Brillouin shift values of the sample (Zhang et al. 2017, Prevedel et al. 2019, Elsayad, Polakova and Gregan 2019b, Elsayad et al. 2019a).

**FIGURE 2:**
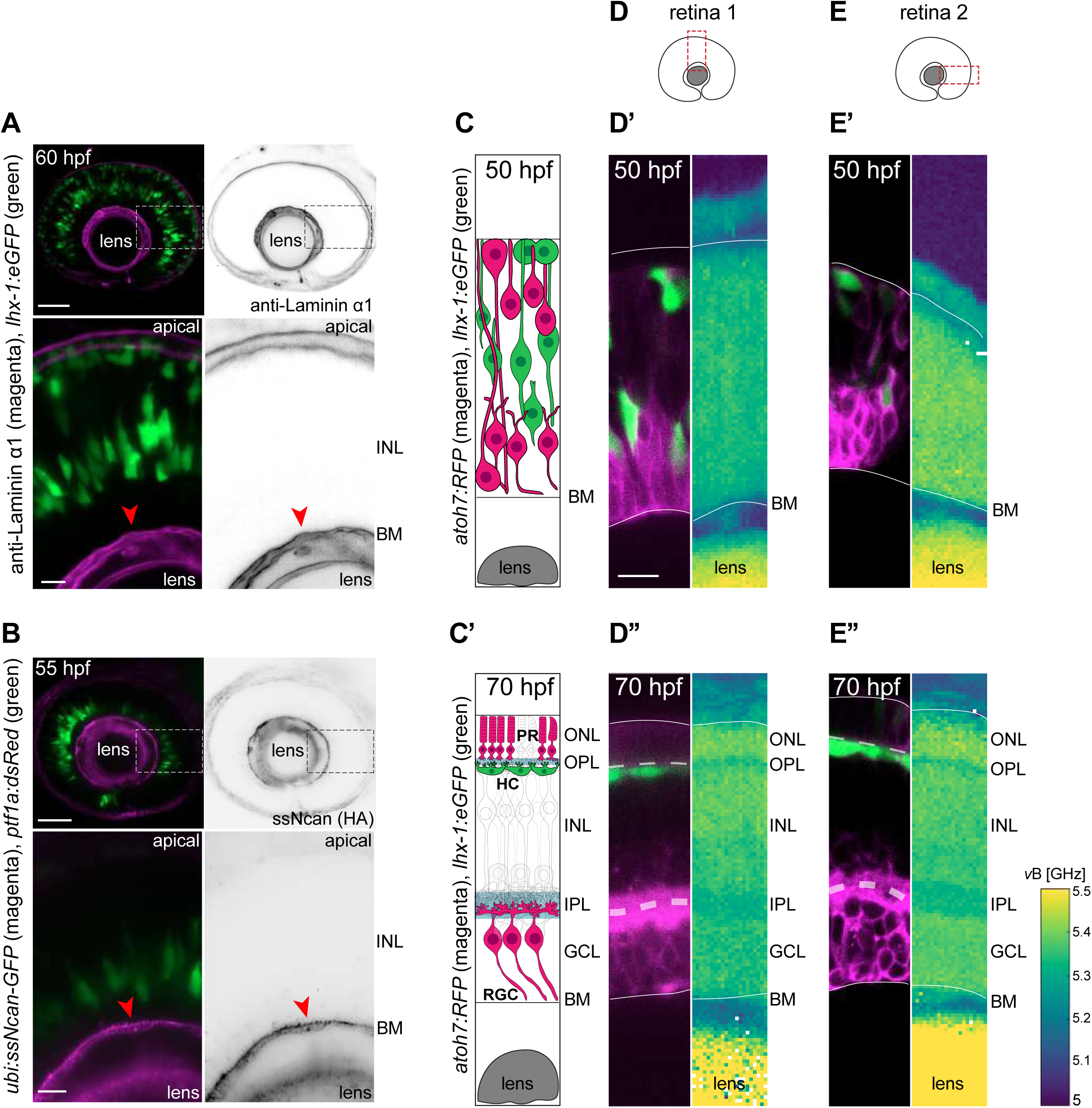
INL is a densely-packed tissue environment which is not dominated by ECM. **(A-B)** INL is not dominated by ECM: **(A)** *Tg(lhx-1:eGFP)* labels HCs [green]. Laminin α1 antibody marks laminin [magenta]. **(B)** *Tg(ptf1a:dsRed)* marks HCs and ACs [green]. *Tg(HA:GFP)* labels hyaluronic acid (HA) [magenta]. Higher magnification insets show enrichment of anti-Laminin α1 and HA in the basement membrane (BM) [red arrowheads]. Scale bars: 50 μm, (insets) 10 μm. **(C-C’)** Scheme of structural organization of the retina during development: **(C)** During HC migration at 50 hpf, **(C’)** After HC layer formation at 70 hpf. HCs [green]. PRs and RGCs [magenta]. **(D-E”)** Brillouin shift maps (right) and their corresponding confocal images (left) of double-transgenic zebrafish of **(D-D’’)** retina 1 and **(E-E’’)** retina 2. Top: 50 hpf, and bottom: 70 hpf. *Tg(lhx-1:eGFP)* labels HCs and ACs [green], *Tg(atoh7:RFP)* is expressed in RGCs and PRs [magenta]. Confocal images were obtained directly after the Brillouin shift measurements. The corresponding Brillouin shift maps of the nuclear layers (ONL, INL, GCL) show a higher Brillouin shift than the plexiform layers (OPL, IPL). Red dashed boxes: imaged region; Lines: apical surface (top), BM (bottom); Dashed lines: OPL (top), IPL (bottom). Scale bar: 10 µm.

Using a custom-built Brillouin microscopy setup (Schlussler et al. 2018), we profiled the *in vivo* Brillouin maps of distinct regions of the zebrafish retina **(Fig 2D, E)**, at different developmental stages: 1) during HC migration (50 hpf) **(Fig 2C)**, and 2) post-HC layer formation (70 hpf) **(Fig 2C’)**, in parallel with confocal fluorescence microscopy.

The Brillouin shift maps revealed that in comparison to the retina neuroepithelium, the lens had higher Brillouin shift at 50 hpf **(Fig 2D’-E’)** which further increased as development progressed (70 hpf) **(Fig 2D’’-E’’)**. At 70 hpf, the retina displays a layered organization composed of five different layers from apical to basal: ONL, OPL, INL, IPL, and GCL **(Fig 2C’)**. Notably, Brillouin shift maps of 70 hpf retinae revealed a layered, heterogenous pattern that matched the retinal layers observed in confocal images **(Fig 2D’’-E’’)**. While the two nuclear-free plexiform layers (OPL and IPL) showed lower Brillouin shifts, the three nuclear layers (ONL, INL, GCL) displayed comparably higher Brillouin shifts. This indicated that a correlation between Brillouin shift maps and the characteristic features of each retinal layer exists, and that Brillouin shift values are influenced by nuclear occupation, as shown previously for fibroblasts (Zhang et al. 2017). However, Brillouin shift values in the INL showed no obvious differences along the apico-basal axis, during **(Fig 2D’, E’)** and after HC migration **(Fig 2D’’, E’’)**. Thus, HC migration seems to occur in a homogenously compressible environment without notable local compressibility gradients.

As Brillouin shift values were higher in regions with high cell-body density compared to the nuclei-free plexiform layers, we tested INL packing at stages of HC peak of migration (50-65 hpf). To this end, we performed live-imaging of double-transgenic zebrafish embryos *Tg(lhx-1:eGFP)* x *Tg(βactin:mKate2-ras)* labeling HCs and membrane of all retinal cells, respectively. No extracellular space between neighboring cells within the INL was detected during periods of HC migration **(Sup-Fig 2A-A’)**. This notion was further supported by confocal imaging of membrane (*Tg(βactin:mKate2-ras)*) and nuclei envelope (*Tg(βactin:eGFP-lap2b)* of all retinal cells, showing that no space was detected in between the neighboring nuclei in INL **(Sup-Fig 2B-B’)**. Thus similar to the neuroepithelial stage (Matejcic et al. 2018), also during neurogenesis, the developing INL features a dense packing of cells and nuclei, implying that HCs migrate in an environment wherein space is limited.

### MIGRATING HORIZONTAL CELLS UNDERGO CELL AND NUCLEAR DEFORMATIONS

We showed that migrating HCs navigate within a crowded environment with neither cellular nor ECM scaffolding structures **(Fig 1C, E, Fig 2A-B),** or compressibility gradients **(Fig 2D-E’’)**. Consequently, HCs are left with two major strategies employed by cells migrating in crowded environments with physical constraints: 1) active generation of migratory tracks by protease-dependent local ECM degradation or remodeling (e.g. used by mesenchymal cells such as fibroblasts during wound healing, and aggressively migrating tumor cells) (Wolf et al. 2007, Wolf and Friedl 2011, Krause and Wolf 2015), or 2) amoeboid adaptations of their shape, migration path and direction to the limited available space (e.g. seen for leukocytes like neutrophils and dendritic cells) (Lammermann and Sixt 2009, Friedl and Wolf 2010).

Given that no major ECM components were detected in the INL **(Fig 2A-B)**, protease-dependent strategies were unlikely to be of major importance in this context. To underline this notion, tissue integrity was probed using double-transgenic zebrafish retinae *Tg(lhx-1:eGFP)* x *Tg(bactin:mKate2-ras)* labeling HCs and PRs, and membrane of all retinal cells, respectively. Consistent with lack of ECM in the INL, we did not detect any tissue ruptures or holes, neither at the peak of HC migration nor after HC migration using time-lapse imaging and immunostainings **(Sup-Fig 2A-A’)**. Thus, we considered it unlikely that HCs employ a protease-dependent path generation strategy to move through the crowded retina.

Typically, amoeboid migrating cells undergo cytoplasmic and nuclear deformations which allow them to navigate through crowded environments either *in vivo* (Friedl, Wolf and Lammerding 2011, Wolf et al. 2003, Salvermoser et al. 2018, Manley et al. 2020) or in narrow channels *in vitro*. We thus asked whether such deformations accompanied and/or influenced HC migration behavior. To achieve mosaic labeling of cells, we either used blastomere transplantation of *Tg(lhx-1:eGFP)* to mark cell bodies, or injected trbeta2:tdTomato and LAP2b:eGFP DNA constructs to visualize cell bodies and nuclear envelopes of HCs, respectively. We observed that migrating HCs exhibited a wide variety of cellular and nuclear shape changes, ranging from elongated to bended **(Fig 3A-A’’)**.

**FIGURE 3:**
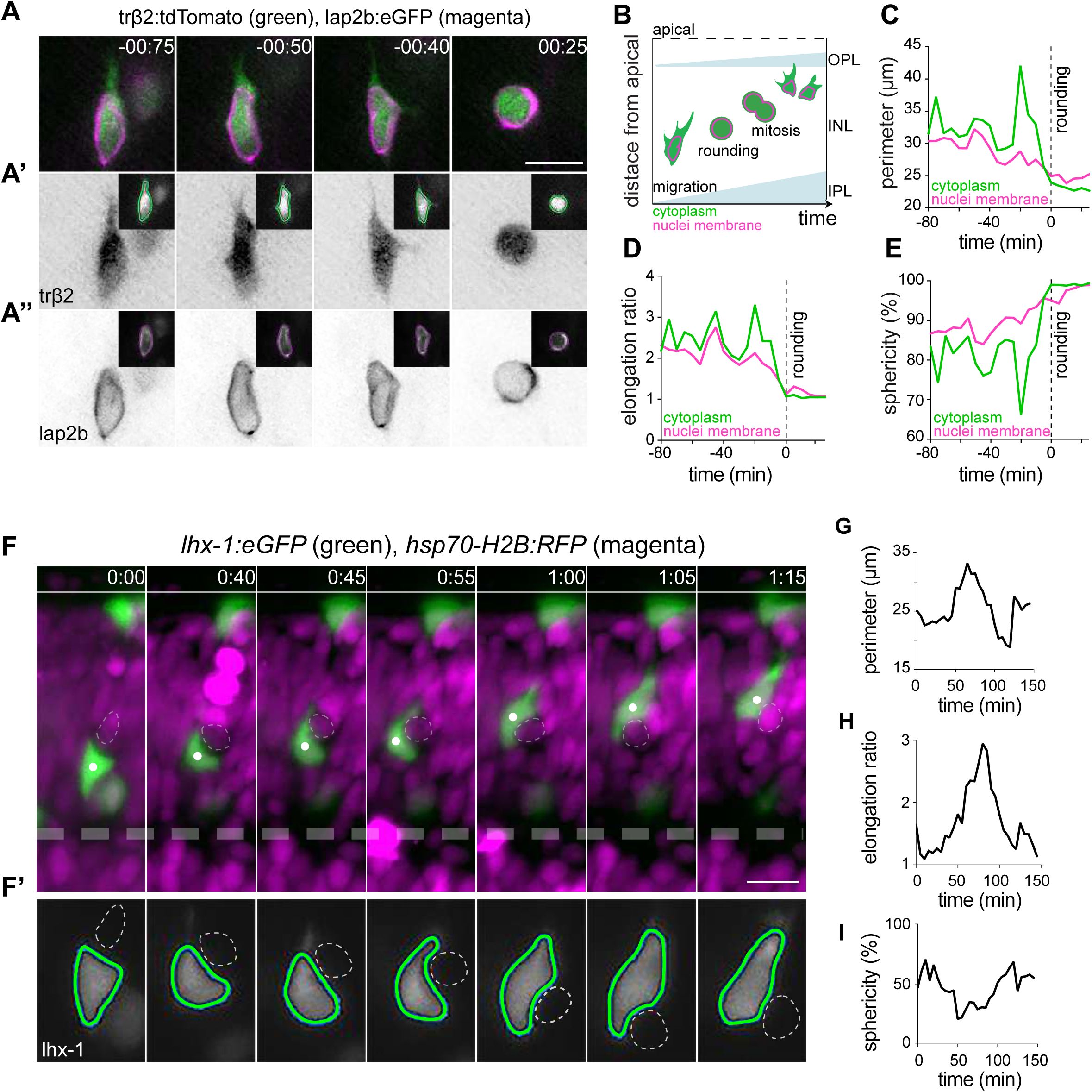
HCs undergo cell and nuclear deformations as they migrate through the crowded INL. **(A-E)** The cell bodies and nuclei of migrating HCs are deformable. **(A)** Time-lapse of an HC during migration and at entry into mitosis. The HC cell body is visualized by *trβ2:tdTomato* [green], and its nuclear envelope by *lap2b:eGFP* [magenta] DNA constructs. Insets show the corresponding segmented contours of the tracked HC’s **(A’)** cell body (green lines), and **(A’’)** nucleus (magenta lines) to extract cellular and nuclear morphodynamic features in (C-E). Scale bar: 10 µm. **(B)** Schematic representation of a typical basal-to-apical migration trajectory of an HC which undergoes mitosis en route. During mitosis, HCs switch from an elongated shape into a spherical morphology. **(C-E)** Graphs show quantification of the dynamics of the cell and nucleus shape-changes of the migrating HC depicted in (A): **(C)** Perimeter (μm), **(D)** elongation ratio, and **(E)** sphericity (%). Note that the minimal perimeter (μm), minimal elongation ratio and maximal sphericity (%) are reached upon cell rounding during mitosis. Dashed lines: onset of HC rounding. **(F-I)** Migrating HCs squeeze in the crowded retina to overcome local physical obstacles. **(F)** Stills from light-sheet time-lapse imaging show that the migrating HC [green] undergoes cell-shape deformations to circumvent the rounded mitotic nuclei of its neighboring cell (dashed circle). *Tg(lhx-1:eGFP)* labels HCs [green], *Tg(hsp70-H2B:RFP)* marks nuclei of all retinal cells [magenta]. White dot: tracked HC; line: apical surface; Dashed line: IPL. Scale bar: 20 µm. **(F’)** The automated segmented contours of the cell body of the migrating HC [green line] and the nucleus of its neighboring cell (dashed circle) from (F). **(G-I)** Graphs represent quantifications of cell morphodynamic changes in HC from (F-F’): **(G)** perimeter (μm), **(H)** elongation ratio, and **(I)** sphericity (%). Time in h:min (A, F).

To quantify these morphological alterations, we used the open-source Icy platform (de Chaumont et al. 2012) (http://icy.bioimageanalysis.org) for automated segmentation of HC cell and nucleus contours (Manich et al, 2020) **(Fig 3A’-A’’, Video 4)**. We measured perimeter (μm), elongation ratio and sphericity (%) of cell body and nucleus of HCs during migration, as well as during their final mitosis **(Fig 3B)**, on their basal-to-apical journey. At mitosis, HCs displayed an elongation ratio of ∼1 **(Fig 3D)** and a sphericity of ∼100% **(Fig 3E)**. This implied that cell bodies and nuclei of mitotic HCs adopt a spherical shape **(Fig 3A’-A’’)**, as seen for most animal cells entering mitosis (Taubenberger, Baum and Matthews 2020), and validated our method. During migration, HCs exhibited a more elongated cellular morphology and underwent multiple cellular deformations **(Fig 3A’, Sup-Fig 3A-A’’)** as was confirmed by frequent changes in their perimeter (μm), elongation ratio and sphericity (%) **(Fig 3C-E, Sup-Fig 3B-D, Video 4)**. These cell shape changes were accompanied by alterations in nuclear perimeter (μm), elongation ratio and sphericity (%) **(Fig 3A’’, C-E)** and dynamic indentations of the nucleus **(Sup-Fig 3E-E’, Video 5)**.

To test whether a correlation between encountered tissue obstacles and shape changes of HCs existed, we monitored HC migration (*Tg(lhx-1:eGFP)*) in relation to the surrounding local environment (*Tg(hsp70-H2B:RFP)*). We noted that migrating HCs featured cellular deformations when encountering a neighboring cell that entered mitosis **(Fig 3F-I, Video 6)**. In other cases, when the tissue seemed impassable, HCs changed both their direction of migration and cellular shape, often taking less direct routes to their final position. Notably, the adaptive cellular and nuclear deformations shown by HCs were reversible after the physical barrier was circumvented **(Fig 3F-I)**. Together, these data support the notion that migrating HCs tailor their cellular and nuclear shapes, paths and directionalities to their surrounding tissue environment. This behavior does not generate migration trails but could rather serve as a space-adaptation strategy enabling HCs to successfully move within the densely-packed and dynamically complex retina. Many types of amoeboid migration from *Dictyostelium* to leukocytes (Friedl and Wolf 2010, Trepat, Chen and Jacobson 2012, Arts et al. 2021) show similar morphological changes. Therefore, our results suggest that HCs undergo amoeboid-like migration in the zebrafish retina.

### HORIZONTAL CELLS DISPLAY A POLARIZED FRONT-REAR MORPHOLOGY

To examine whether HC migration displayed further cell biological characteristics of amoeboid migration, we investigated their protrusion activity. Classically, two types of protrusions can influence amoeboid migration: 1) amoeboid bleb-guided migration (e.g. seen in macrophages) (Yoshida and Soldati 2006, Charras and Paluch 2008, Bergert et al. 2012), 2) amoeboid pseudopodal-guided migration (e.g. used by *Dictyostelium* and neutrophils) (Trepat et al. 2012, Friedl and Wolf 2010). To elucidate HC protrusion activity, we monitored HC membranes in *Tg(ptf1a:Gal4-VP-16,UAS:gap-YFP)* embryos. As seen for many other cell types (Boss 1955, Burton and Taylor 1997, Fishkind, Cao and Wang 1991, Boucrot and Kirchhausen 2007), HCs displayed membrane blebs only during mitosis and never during migration **(Sup-Fig 4A, Video 7)**.

In contrast, we found that HCs feature multiple dynamic protrusions with different directionality as they moved sideways (tangentially), up (towards the apical) or down (towards the basal) within the INL. These protrusions showed different thicknesses, lengths, morphologies (branched vs. unbranched) and orientations, and were dynamically extended and retracted from the cell soma in multiple directions **(Fig 4A, Video 7)**. At times, two or multiple protrusions were seen to extend from the cell body at the cell front **(Fig 4A)**, another feature also reported in amoeboid migrating cells (Weber et al. 2013, Renkawitz et al. 2019, Kameritsch and Renkawitz 2020).

**FIGURE 4:**
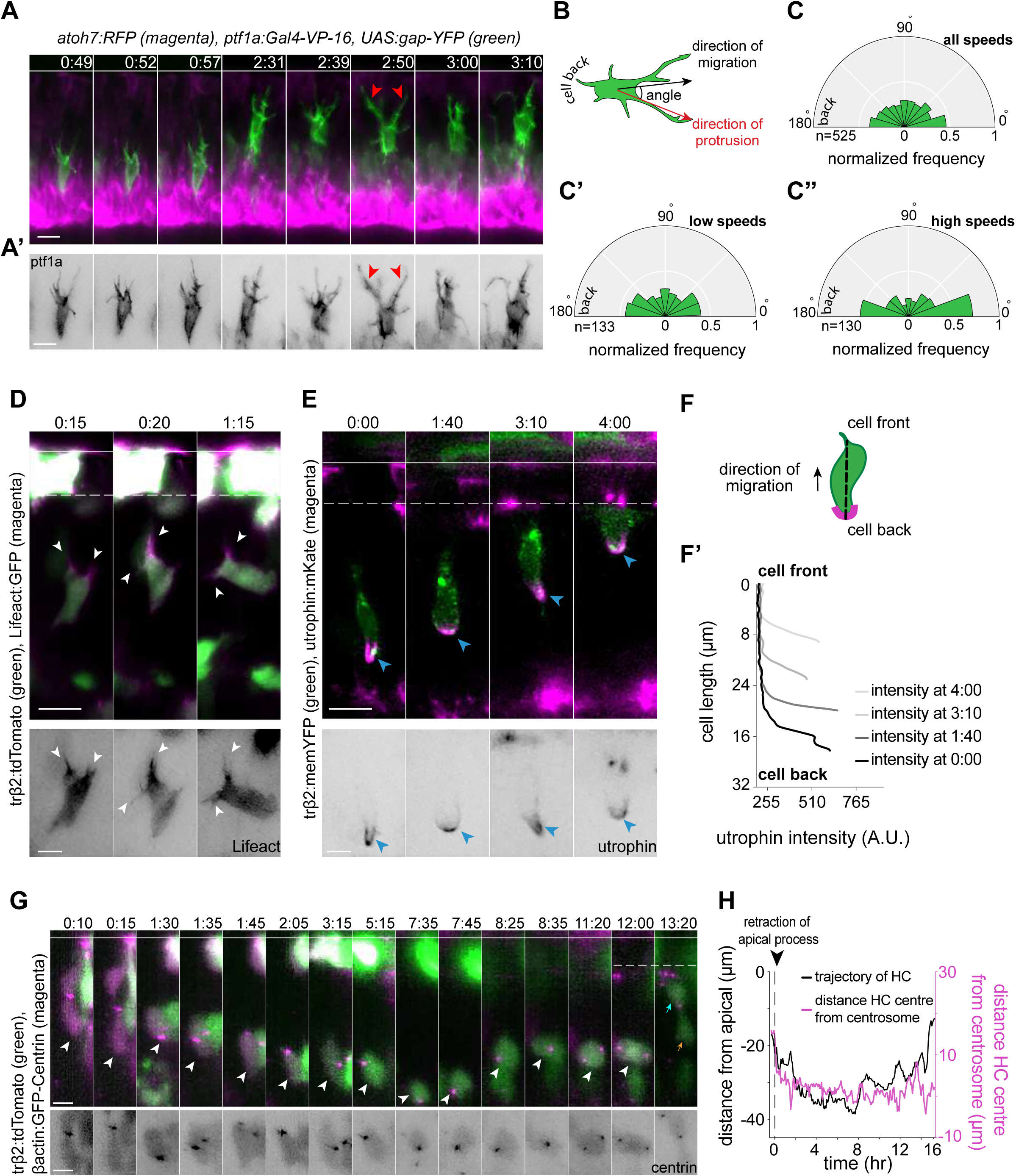
HCs feature hallmarks of protrusion-based amoeboid-like migration. **(A)** Time-lapse sequence of highly dynamic membrane protrusions pointing at different directions (red arrowheads) in a migrating HC. *Tg(ptf1a:Gal4-VP-16,UAS:gap-YFP)* labels membrane of HCs [green], *Tg(atoh7:RFP)* marks RGCs and PRs. Time interval = 1 min. Scale bar: 50 μm. **(A’)** Higher magnification insets of HC cell membrane from (A). Scale bar: 5 μm. **(B)** Scheme of protrusion angles measurements. **(C-C’’)** The frequency distribution of the angle between the direction of instantaneous movement of HC and its protrusions; **(C)** throughout the tracked basal-to-apical journey (t=4hr), **(C’)** at low speeds: speeds below 25 percentile of that cell speed (< 0.38 μm/min), **(C’’)** at high speeds: above 75 percentile of this cell’s speed (> 0.93 μm/min). Minimum and maximum speeds observed for this cell are 0.06 µm/min and 4.44 µm/min respectively. A protrusion pointing exactly toward the direction of cell movement has an angle of 0°, and a protrusion pointing exactly opposite has an angle of 180°. The radius indicates the normalized frequency for each angle bin, i.e. the number of frames observed with the angle belonging to the particular angle bin for the given velocity condition normalized by the total number of frames observed in the given velocity condition. All measurements are from the migrating HC in (A-A’). **(D-F’)** F-actin distribution in migrating HCs: **(D)** trβ2:tdTomato and Lifeact-GFP DNA constructs mark HCs [green] and all filamentous F-actin [magenta], respectively. Lifeact-GFP is detected in the leading protrusions (white arrowheads) and the cell cortex. Scale bar: 20 μm. Bottom: close-up of Lifeact-GFP [gray] from (D). Scale bar: 5 μm. **(E)** trβ2:memYFP is expressed in HCs [green] and utrophin-mKate marks the stable filamentous F-actin [magenta]. Utrophin is enriched at the cell back (blue arrowheads). Scale bar: 20 μm. Bottom: high magnification of utrophin:mKate [gray] in migrating HC from (E). Scale bar: 5 μm. **(F)** Scheme of line scan measurements of utrophin:mKate fluorescence intensity. **(F’)** Average utrophin:mKate fluorescent intensity profile of images at (E). Cell front and back are determined by the direction of HC movement. **(G-H)** Time-series showing the dynamics of centrosome position in HCs from birth to final positioning. **(G)** bactin:GFP-Centrin and trb2:tdTomato DNA plasmids label centrosomes [magenta] and HCs [green], respectively. White arrowhead: HC. Cyan and orange arrows: sister HCs after mitosis (t=12:00). Bottom panel shows close-up of bactin:GFP-Centrin [gray] in migrating HC. White line: ONL; Dashed white line: HC layer. Scale bar: 20 μm. **(H)** Graph showing migration trajectory of the represented HC (black line) and the distance between its centrosome and center (magenta line) throughout HC migration. Arrowhead: time of detachment from the apical process. Time in h:min (A-G).

To analyze the overall orientation of protrusions with respect to the direction of HC movement, tips of each protrusion were manually tracked in individual HCs (n=6) **(Fig 4A-C’, Sup-Fig 4F-I’’’, Video 8)**. The angle between each protrusion and direction of instantaneous cell movement was then quantified **(Fig 4B)** throughout the tracked basal-apical migration of each HC. This analysis revealed that the overall probability of finding protrusions with orientations parallel to the direction of cell movement is higher than finding protrusions pointing in other directions **(Fig 4C, Sup-Fig 4F’-I’)**, implying that HCs show some type of front-back polarity while migrating. Moreover, we observed that the protrusions were typically more randomly oriented when HCs moved with lower cell speed (cell speeds below 25 percentile of the measured speeds of that cell during its tracked trajectory) **(Fig 4C’, Sup-Fig 4F’’-I’’)**. In contrast, when cells moved with high speed (cell speeds above 75 percentile of the measured speeds of that cell during its tracked trajectory), the frequency of protrusions with 0° (cell-front) and 180° (cell-back) was overall higher **(Fig 4C’’, Sup-Fig 4F’’’-I’’’)**.

One possible component that could be responsible for front-rear polarity in HCs during migration is the asymmetric distribution of stable F-actin as reported for amoeboid migrating cells including leukocytes (Cassimeris, McNeill and Zigmond 1990), dendritic cells in 3D (Insall and Machesky 2009, Lammermann et al. 2008), neutrophils (Yoo et al. 2010, Manley et al. 2020, Barros-Becker et al. 2017) and neutrophil-like cells *in vitro* (Cooper, Bennin and Huttenlocher 2008). We thus investigated F-actin distribution in HCs using two distinct actin bioprobes: Lifeact (17 amino acids of yeast Abp140), which labels all filamentous F-actin structures (Yoo et al. 2010, Fritz-Laylin et al. 2017), and utrophin (calponin homology domain) (Utr-CH) that has been shown to preferentially bind to a more stable cortical population of F-actin (Burkel, von Dassow and Bement 2007, Riedl et al. 2008, Yoo et al. 2010, Belin, Goins and Mullins 2014, Barros-Becker et al. 2017). We found that while Lifeact was observed in the cell soma and protrusions **(Fig 4D, Video 9)**, Utr-CH was absent from the cell soma, membrane-proximal regions and protrusions **(Fig 4E, Video 10)**. Instead, measurements of Utr-CH fluorescent intensity profiles along the cell axis showed its enrichment at the back of the migrating HC **(Fig 4D, Fig 4F’, Sup-Fig 4E)**. This is akin to the structure seen at the cell-rear of amoeboid migrating cells also known as uropod (Barros-Becker et al. 2017, Manley et al. 2020, Hind, Vincent and Huttenlocher 2016). Monitoring Utr-CH during HC migration showed that the uropod-like structure underwent dynamic changes of extension and retraction **(Sup-Fig 4C-D, Video 10)**, a feature previously reported for amoeboid migrating neutrophils (Manley et al. 2020). Front-back polarity manifestation of migrating cells is typically accompanied by asymmetric positioning of organelles, including centrosomes (Kupfer, Louvard and Singer 1982, Luxton and Gundersen 2011). We thus monitored distribution and dynamics of centrosomes during HC migration, using Centrin:GFP (centrosome marker) and *trβ2:tdTomato* (HC marker) DNA injections **(Fig 4G, Sup-Fig 5A-D)**. Our analysis revealed that HC centrosomes displayed a highly variable and dynamic localization and continuously shifted their positions from the cell-front to the cell-back while occasionally staying in the cell’s middle **(Fig 4G-H, Sup-Fig 5A)**. This oscillating configuration suggested that centrosome position did not directly influence the direction of HC movement. Overall, we concluded that similar to amoeboid moving cells, HCs also acquire a polarized morphology with persistent rearward polarization of stable F-actin, most likely without contribution of centrosome position. This further supports the idea that HCs employ amoeboid-like migration strategies to move forward in the zebrafish retina.

### OVEREXPRESSION OF LAMIN A IMPACTS THE EFFICIENCY OF HORIZONTAL CELL MIGRATION AND LAMINATION

We showed that HCs undergo frequent cellular and nuclear deformations **(Fig 3A-D)** while migrating through the densely-packed INL **(Sup-Fig 2A-B’’)**. Consequently, we asked whether changing the properties of nuclei as the biggest and bulkiest cell organelle (Martins et al. 2012, Lammerding 2011), could impact HC migration. Some nuclear properties are determined by the differential expression of type V intermediate filament proteins of A- and B-type lamins which are part of the nuclear lamina (Broers et al. 1997, Gruenbaum et al. 2005, Lammerding et al. 2006, Gerace and Huber 2012, Burke and Stewart 2013). Particularly A-type Lamins (A, C and C2) are inversely correlated to the deformability of the nucleus as increasing their expression levels has been linked to decreasing nuclear deformability (Lammerding et al. 2006, Harada et al. 2014, Rowat et al. 2013, Swift et al. 2013, McGregor, Hsia and Lammerding 2016). It was shown that nuclei in the developing zebrafish retina only express negligible levels of A-type lamins (Yanakieva et al. 2019). Thus, we probed whether and how changing nuclear properties by increasing the expression of Lamin A (LMNA) at the tissue-scale could influence HC migration.

To test this, we generated a zebrafish transgenic line *Tg(hsp70:LMNA-mKate2)* in which LMNA overexpression in all cells is induced upon heat-shock. We then quantitatively analyzed HC migration and layer formation in this condition. First, we studied HC layer formation 40 hrs after heat-shock, in fixed samples of *Tg(lhx-1:eGFP)* (as controls) **(Fig 5A)** and *Tg(lhx-1:eGFP)* x *Tg(hsp70:LMNA-mKate2)* double transgene retinae **(Fig 5B)**, at 90 hpf, a developmental stage at which HC layer formation is complete in wild-type embryos. In control heat-shocked retinae, all HCs were positioned within the HC layer at this stage **(Fig 5A)**, showing that heat-shock treatment of the embryos does not impair proper HC layer formation. In contrast, in the LMNA overexpressing retinae, many HCs were found at ectopic basal positions, mostly within INL and at times even in the GCL **(Fig 5B)**.

**FIGURE 5:**
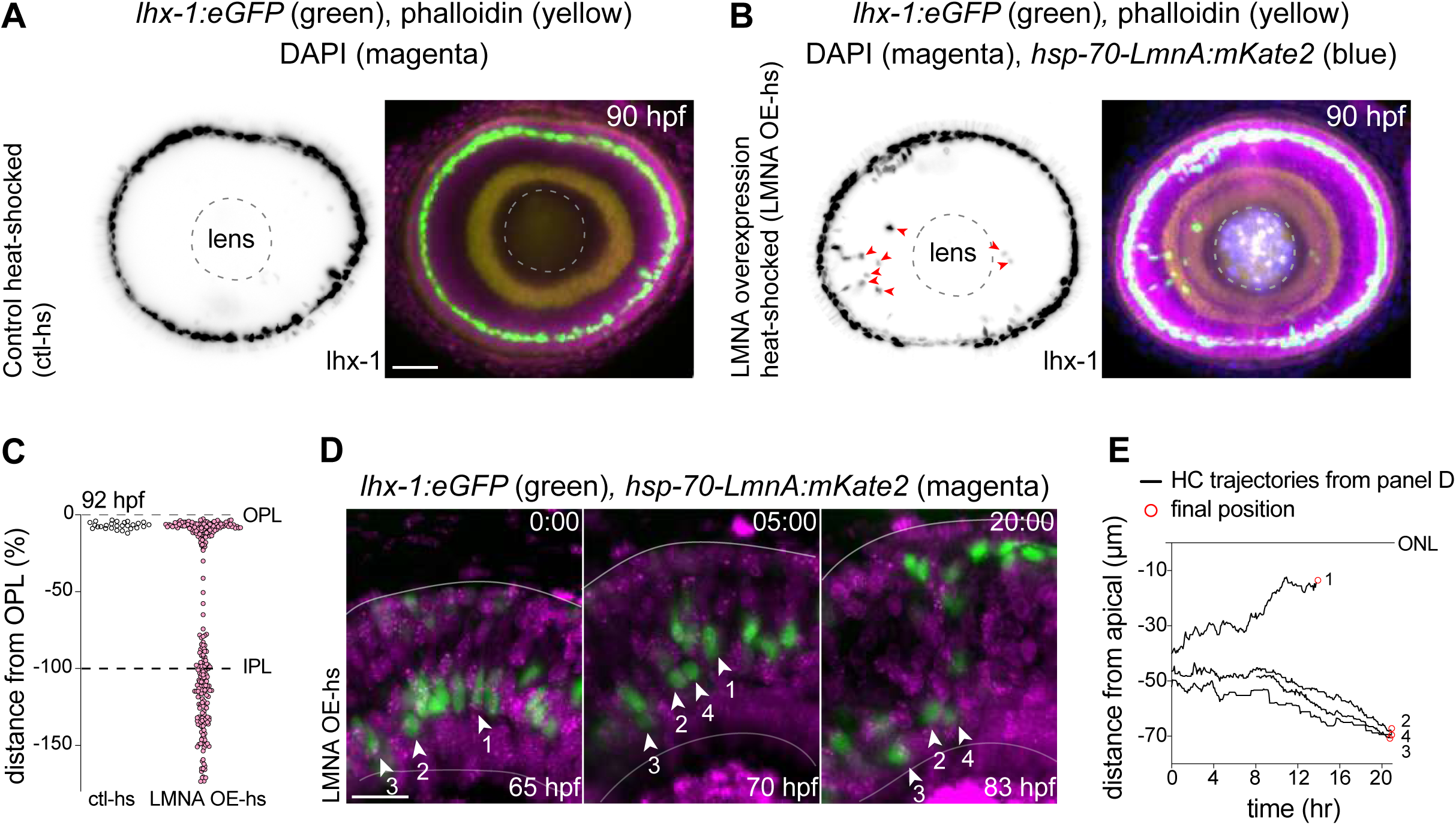
Interference with nuclear properties impairs efficient HC migration and layer formation. **(A-B)** Immunofluorescence images of **(A)** control heat-shocked (control-hs), and **(B)** (LMNA overexpressing heat-shocked (LMNA OE-hs) at 90hpf. *Tg(lhx-1:eGFP)* embryos were used for ctl-hs, and *Tg(lhx-1:eGFP) x Tg(hsp70:LMNA-mKate2)* double-transgene retinae for LMNA OE-hs experiments. *Tg(lhx-1:eGFP)* marks HCs [green], *Tg(hsp70:LMNA-mKate2)* is expressed in nuclear envelopes upon heat-shock [blue], DAPI labels nuclei [magenta], and phalloidin marks actin [yellow]. Red arrowheads: ectopically positioned HCs, dashed circle: lens. Scale bar: 50 μm. **(C)** Position of HCs relative to OPL in control heat-shocked (ctl-hs) (n=27, N=5), and LMNA overexpressing heat-shocked retinae (LMNA OE-hs) (n=283, N=11) at 92 hpf. **(D)** Stills from a 20hr time-lapse of *Tg(lhx-1:eGFP) x Tg(hsp70:LMNA-mKate2))* double-transgene retinae after heat-shock (LMNA OE-hs). Arrowheads: tracked HCs. Scale bar: 20 μm. **(E)** Migration trajectories of tracked HCs from (D). HC n-2, n-3 and n-4 remain ectopically positioned in the basal part of the retina. Time in h:min (D).

To assess whether the apical migration of the ectopically-positioned HCs is delayed or abrogated as a result of LMNA overexpression, we performed long-term *in vivo* imaging (40-45 hrs) of *Tg(lhx-1:eGFP)* x *Tg(hsp70:LMNA-mKate2)* double transgene embryos and monitored the overall HC layer formation until animals reached 92 hpf. We found that while the majority of HCs in LMNA overexpressed condition reached the HC layer at 92 hpf, a subset of HCs remained at ectopic basal positions **(Fig 5C)**. Analyzing the migration trajectories of tracked HCs revealed that in the LMNA overexpressing condition, many HCs failed to move back toward the apical INL **(Fig 5D-E)** and consequently resided in ectopic basal positions away from their final position **(Sup-Fig 5E)**. This implies that changing the nuclear laminar composition at the tissue-scale hampers the migration efficiency of HCs and thereby perturbs HC layer formation.

### INNER PLEXIFORM LAYER ACTS AS A BARRIER FOR HORIZONTAL CELL MIGRATION

Since HCs are unlikely to remodel their environment to generate their path, we wondered whether their migration path could be impacted by physical barriers in the tissue. In the developing zebrafish retina, five layers with discrete properties **(Fig 2C-E’’, Sup Fig 2B-B’’)** and topographical features emerge during neuronal lamination. We previously showed that IPL, once it is formed, negatively influences the depth of HC migration, as HCs did not basally pass it (Amini et al. 2019). This suggested that IPL, despite being devoid of cell-bodies may act as a non-passable obstacle for migrating HCs. This is most likely due to the intermingled axonal terminals of BCs, dendritic trees of ACs and RGCs, and MG processes which form a dense neuropil enriched with membrane **(Sup Fig 2B-B’’, Sup Fig 5F)**.

To test this hypothesis, we set out to drive HCs to positions below IPL before its formation and explore how HCs deal with IPL once it was formed. To this end, we genetically eliminated RGCs, using a validated *atoh7* morpholino (Kay et al. 2004, Kay et al. 2001, Pittman, Law and Chien 2008) that results in RGC depletion, and thereby delays IPL formation. In this condition, ACs are found at the most basal layer (Kay et al. 2001) intermixed with occasional HCs (Weber et al. 2014). Our immunofluorescence stainings of *Tg(lhx-1:eGFP)* retinae revealed that in *atoh7* morphants at 50-55 hpf, the maximal depth of HC migration increased significantly and that many HCs were located adjacent to the basement membrane at the most basal side of the retina, a depth never seen for HCs in control embryos **(Fig 6A, C)**. Thus, upon interference with RGC emergence, HCs reached deeper positions in the tissue by moving beyond their typical basal stopping-point. This suggests that the RGC layer acts as a basal brake for HCs during their apical-to-basal journey.

**FIGURE 6:**
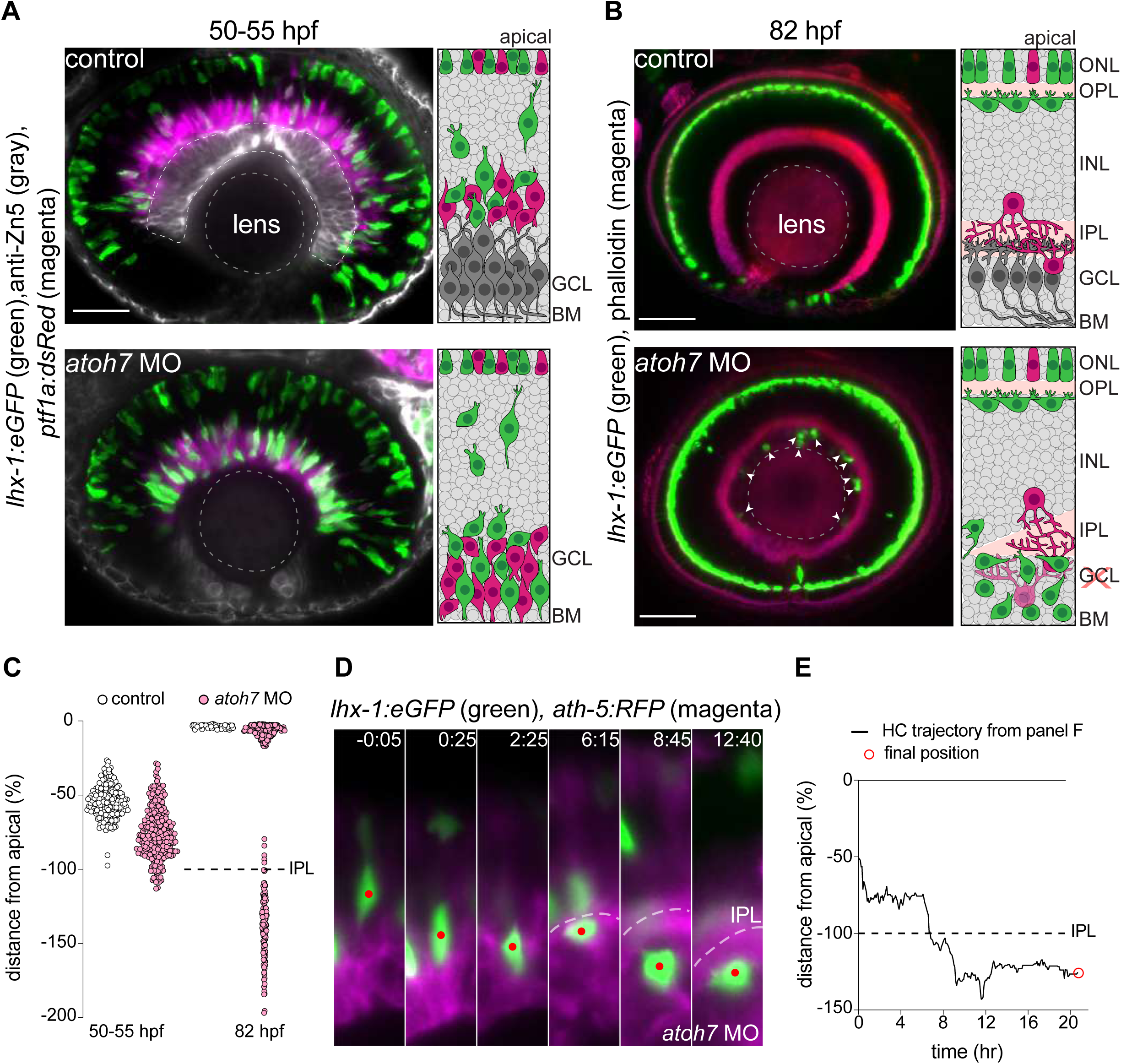
IPL POSES A BARRIER FOR HORIZONTAL CELL MIGRATION. **(A-B)** Immunofluorescence images of control (ctl) (top) and *atoh7* morpholino (*atoh7* MO) (bottom). Schematic representations of each condition is on the right. **(A)** Control and *atoh7* MO retinae at 50-55 hpf retinae. *Tg(lhx-1:eGFP)* labels HCs and PRs [green], *Tg(ptf1a1:dsRed)* is expressed in ACs and HCs [magenta], anti-Zn5 marks RGCs [gray]. In *atoh7* MO retinae, the most basal layer is devoid of GCL [gray in ctl] and is instead filled with ACs and HCs [magenta, green]. Dashed circle: lens. Scale bar: 50 μm. **(B)** Control and *atoh7* MO retinae at 82 hpf. *Tg(lhx-1:eGFP)* labels HCs [green], phalloidin marks actin [magenta]. In *atoh7* MO retinae, many HCs remains trapped underneath IPL, formation of which is delayed. Arrowheads: trapped HCs; dashed circle: lens. Scale bar: 50 μm. **(C)** Position of HCs relative to the apical surface in control (n=196, N=12) and *atoh7* MO (n=261, N=14) retinae at 50-55 hpf, and in control (n=76, N=11), and *atoh7* MO (n=312, N=15) at 82 hpf. Dashed line: IPL. **(D)** Time-series of a tracked HC (red dot) in *atoh7* MO retinae. Red dot: tracked HC; dashed line: IPL. Scale bar: 20 μm. **(E)** Migration trajectory of the tracked HCs from (D). Time in h:min (D-E).

Because these basal locations are completely different from INL wherein HC migration occurs, we asked whether ectopically located HCs are able to reach their apical layer beneath OPL. To determine this, we monitored HC migration in *atoh7* morphant retinae until 82 hpf. We found that before IPL formation, HCs successfully turned back apically and reached the HC layer in *atoh7* morphants, implying that the ability of HCs to move back apically is not abrogated in the absence of RGCs. In contrast, during or after IPL formation, HCs failed to reach the HC layer **(Fig 6D-E)**. Consequently, while all HCs reached the HC layer at 82 hpf in control retinae, many remained ectopically constrained at positions below IPL, in RGC depleted conditions **(Fig 6B,C)**. While, these ectopically located HCs were still able to move in all dimensions (up, down, lateral) below IPL, they failed to migrate apically through it. As a result, HCs remained trapped below IPL until they underwent apoptosis, evidenced by their progressive fragmentations, immobility and ultimately disappearance. Overall, we conclude that IPL despite being more compressible than nuclear layers **(Fig 2D-E’’)**, represents a barrier through which migrating HCs are not able to pass when trapped beneath. Thus, IPL likely poses a limit to the morphological adaptability of HCs and their nuclei.

## DISCUSSION

We here showed that HCs employ space adaptation strategies to navigate through the complex and crowded environment of the developing retina. In particular, we revealed that HCs repeatedly and reversibly adjust their cellular and nuclear morphology, and direction of movement to the constraints they encounter within their surrounding densely-packed tissue. Because we found that the migratory behavior and morphology of HCs share many hallmarks of amoeboid migration, we refer to HC migration as “amoeboid-like neuronal migration”. To the best of our knowledge, this is the first study describing a neuronal cell type that undergoes amoeboid-like migration in a part of the developing CNS. We further uncovered that changing tissue properties can feedback on the efficiency of ameboid-like neuronal migration and layer formation most likely by influencing the space-negotiation capability of HCs.

As opposed to neocortical neurons including projection neurons which *en route* to their destination switch from multipolar to bipolar morphology and resume unidirectional radial migration along radial glia fibers (Nadarajah et al. 2003, Nadarajah et al. 2001, Cooper 2014), HCs do not seem to travel along radially oriented progenitors, MGs or BCs. During long stretches of their migration, HCs follow unconventional tortuous migratory tracks while frequently alternating between radial (up and down) and tangential (lateral) routes. Such migration trajectories also set HCs apart from other emerging retinal neurons including PRs (Rocha-Martins et al. 2021) and RGCs (Icha et al. 2016a), which display bipolar morphologies and remain constrained to radial routes due to their anchored processes.

That HCs move without anchorage during most of their journey allows them to move freely in all dimensions, a feature which in combination with HCs’ flexible morphodynamic properties assists them to overcome obstacles of moving in the crowded environment by taking stochastic migration tracks (Amini et al. 2019). Thus, while HCs reproducibly and robustly reach the HC layer, their path selection is not intrinsically programed but rather influenced by the cellular surroundings and the tissue-scale parameters encountered in their local environment.

The amoeboid-like migration mode exhibited by HCs is not based on bleb formation but correlates with multiple highly dynamic actin-filled protrusions. The exact nature of these protrusions, and whether they directly drive HC migration, allow HCs to explore potential environmental cues or both, remain to be further explored. Our observation that protrusions are simultaneously extended towards multiple directions, especially during periods in which HCs stay stationary, suggests that they are rather involved in probing the tissue environment and pathfinding than directly propelling the movement. Such exploratory roles have been proposed in amoeboid moving cells including *Dictyostelium* during chemotaxis, leukocytes and neutrophils (Gupton et al. 2005, Wu et al. 2012, Leithner et al. 2016, Vargas et al. 2016, Fritz-Laylin et al. 2017, Gerisch and Hess 1974). Future experiments that specifically interfere with protrusion formation or maintenance will shed light on their exact role in HC migration and layer formation.

We currently do not understand the mechanism(s) and forces that move HCs forward. Many amoeboid migrating cells display a front-rear polarity wherein stable F-actin filaments are asymmetrically enriched at the highly contractile uropod (Hind et al. 2016, Bergert et al. 2012, Lammermann and Sixt 2009). Our finding that migrating HCs display strikingly similar polarized morphology implies that they may also use uropod contraction as a pushing force to move forward. Unraveling the spatiotemporal molecular machineries of cell polarity, force-generation, the signaling and the cytoskeletal elements that drive them will be exciting areas for future studies.

Using *in vivo* Brillouin microscopy, we showed that the Brillouin shift maps of INL remain relatively homogenous throughout HC migration. That no obvious compressibility gradient was observed along the apico-basal axis of INL during HC migration implies that tissue compressibility could have a permissive rather than an instructive role. The fact that interfering with tissue-wide components such as properties of the nuclear lamina of HCs and their surrounding cells, impedes HC migration efficiency and successful layer formation further argues in this direction.

An additional tissue-wide feature that influences HC migration is the emergence of IPL. We previously reported that the depth of HC migration correlates with IPL emergence and that once it is formed, HCs do not pass beyond it on their apical-to-basal journey (Amini et al. 2019). This together with our finding that HCs get trapped beneath IPL in RGC-depleted retinae, strongly suggests that IPL acts as a steric hindrance through which HCs cannot penetrate in either direction. While this interpretation may seem at odds with the finding that IPL is more compressible than INL according to the Brillouin shift profile, it is possible that the fibrillar arrangement of axonal and dendritic processes within IPL poses a net-like obstacle with low porosity that is below the deformation capability of the HC nuclei. This idea is in line with studies that showed that migration efficiency is optimal at pore diameters that match or range slightly below the diameter of cell’s nucleus.

It remains unknown what external cues guide HC migration to ensure that HCs always find their accurate functional position while avoiding entrapment within the crowded retina. Such cues could either come in the form of mechanical gradients or chemical signaling or a combination of both. In the future, it will be important to explore the guidance cues that trigger reorientation of HCs toward the apical side where they later reside and function. It will be important to find their source, to understand how they change in space and time, and how HCs sense, integrate and prioritize these cues within the local structural features of their surroundings to successfully find and reach their destination.

Taken together, this study reveals that in addition to the numerous neuronal migration modes characterized so far, neurons can also undergo amoeboid-like migration in an important part of the developing CNS, the retina. The ability to undergo direction, cell- and nuclear-shape changes allows HCs to evade rather than degrade encountered barriers in the crowded tissue environment. It will be interesting to explore whether this mode of migration is specific to HCs in the zebrafish retina or conserved in retinae of other organisms and/or other regions of the brain. As multipolar migration modes are observed in many parts of the developing CNS, it is likely that amoeboid-like neuronal migration is widespread in diverse systems. Similarities and differences of what influences amoeboid-like migration in different systems will teach us more about the intricate development of the nervous systems in vertebrates of all kinds.

## MATERIALS AND METHODS

### 1. ZEBRAFISH WORK

#### 1.1. Zebrafish husbandry

Wild-type TL zebrafish (*Danio rerio*) and transgenic lines were maintained and bred at 26°C as previously described. Embryos were raised at 28.5°C or 32°C and staged in hours post fertilization (hpf) according to (Kimmel et al. 1995). Embryos were kept in E3 medium, which was renewed daily and supplemented with 0.2 mM 1-phenyl-2-thiourea (PTU) (Sigma-Aldrich) from 8±1 hpf onwards to prevent pigmentation. All animal work was performed in accordance with the European Union (EU) directive 2010/63/EU, as well as the German Animal Welfare act.

#### 1.2. Zebrafish transgenesis

To generate *Tg(hsp70:LMNA-mKate2), a* stable transgenic line containing heat-shock inducible LMNA 1 nl of hsp70:LMNA-mKate2 (Yanakieva et al, 2019) was injected at 36 ng/ul, together with Tol2 transposase RNA at 80 ng/ul in double-distilled (dd)H2O supplemented with 0.05% phenol red (to visualize the injection material) (Sigma-Aldrich) into the cytoplasm of one-cell stage wild-type embryos. F_0_ embryos were raised until adulthood. Germline carriers displaying mKate signal were identified in F_0_ progeny, after heat-shock treatment at 37°C, for 20 min, at 24 hpf. Carriers were then outcrossed with wild-type fish.

#### 1.3. Transgenic lines

Refer to **Table S1** for a list of transgene lines.

#### 1.4. DNA injections

To mosaically label HCs or express proteins of interest in the zebrafish retina, DNA constructs were injected into the cytoplasm of one-cell stage embryos. Constructs were diluted in ddH_2_O supplemented with 0.05% Phenol Red (Sigma-Aldrich). Injected volumes ranged from 1 to 1.5 nl. DNA concentrations were 20-30 ng/µl and did not exceed 45 ng/µl when multiple constructs were injected. See **Table S2** for a list of injected constructs.

#### 1.5. Morpholino experiments

To inhibit RGC formation, *atoh7* morpholino (5’-TTCATGGCTCTTCAAAAAAGTCTCC-3’) (Gene Tools) was injected at 2 ng per embryo into the yolk of one-cell embryos. *p53* morpholino (5’-GCGCCATTGCTTTGCAAGAATTG-3’) (Gene Tools), was co-injected at 2-4 ng per embryo to reduce toxicity and cell death.

#### 1.6. Heat-shock

To induce expression of heat-shock promoter (hsp70)-driven DNA constructs and transgenes except *Tg(hsp70:LMNA-mKate2)*, the Petri dish with 36-42 hpf embryos was placed into a water bath set to 37°C for 30 min or 39°C for 15–20 min. Imaging was started 3–6 h after heat shock. For hsp70-driven LMNA overexpression, *Tg(hsp70:LMNA-mKate2)* embryos were transferred E3-containing 15 ml tubes which were preheated in a water bath for 30 min. Heat-shock started 3-10 h before imaging at 45-48 hpf for 30 min at 39°C, or 15 min at 42°C. Embryos which displayed higher mKate fluorescence intensity were picked for experiments.

#### 1.7. Whole-mount staining of zebrafish embryos

Zebrafish were manually dechorionated and fixed overnight in 4% paraformaldehyde PFA (Sigma-Aldrich) in PBS at 4°C. Embryos were washed 5 times for 15 min with PBS-Triton (PBS-T) 0.8%. Embryos were then permeabilized with 1x Trypsin-EDTA in PBS on ice for different time periods depending on the developmental stage (12 min for 42-50 hpf, 15 min for 56-60 hpf, 20 min for 70-90 hpf). The permeabilization solution was replaced with 0.8% PBS-T and embryos were kept for an additional 30 min on ice before rinsing twice with 0.8% PBS-T. Embryos were then blocked in 10% goat or donkey serum (blocking serum) in 0.8% PBS-T for 3 h at room temperature and were subsequently incubated with primary antibodies diluted in 1% blocking serum in 0.8% PBS-T for 3 days at 4°C. They were then washed 5 times for 30 min with 0.8% PBS-T. After blocking, embryos were incubated with appropriate secondary antibodies and DAPI in 1% blocking serum in 0.8% PBS-T for 3 days at 4°C. Embryos were washed 4 times for 15 min with 0.8% PBS-T before storage in PBS at 4°C until imaging. Refer to **Table S3** for antibodies used.

#### 1.8. Blastomere transplantations

Transplantation dishes were prepared by floating a plastic template in a Petri dish that was half-filled with 1% low-melting-point agarose in E3. Once the agarose solidified, plastic templates were gently removed, leaving an agar mould that contained rows of wells to hold embryos. Embryos at stages high to sphere were dechorionated in pronase (Roche) and dissolved in Danieu’s buffer. Dechorionated embryos were transferred to wells in agarose molds using a wide-bore fire-polished glass pipet. Approximately at the 1000-cell stage, cells from the donor embryos were transplanted into the animal pole of the acceptor embryos using a Hamilton syringe. Transplanted embryos were kept on agarose for about 3-5 h and then transferred onto glass dishes that contained E3 medium supplemented with 0.003% PTU and antibiotics (100 U of penicillin and streptomycin, Thermo Fisher Scientific). Transplanted embryos were identified via fluorescence and imaged from 42 hpf for 24-30 hrs.

### 2. IMAGE ACQUISITION

#### 2.1. *in vivo* light sheet fluorescent imaging

Imaging was performed on a Zeiss Light sheet Z.1 microscope as previously described (Icha et al. 2016b). The system was operated by the ZEN 2014 software (black edition). Briefly, embryos were manually dechorionated and mounted in glass capillaries in 0.9% low-melting-point agarose (in E3) supplemented with 240 μg/mL of tricane methanesulfonate (MS-222; Sigma-Aldrich). The sample chamber was filled with E3 medium containing 0.01% MS-222 and 0.2 mM PTU (Sigma-Aldrich) and was kept at 28.5°C. *Z*-stacks spanning the entire width of the retinal neuroepithelium (90-100 μm, depending on the developmental stage) were recorded with 1 μm optical sectioning every 5 min for 10-40 h with a Zeiss Plan-Apochromat 20x water-dipping objective (Carl Zeiss Microscopy; NA 1.0) and two Edge 5.5 sCMOS cameras (PCO), using double-sided illumination mode. *Z*-stacks were recorded every 5 min for 24-40 h, with double-sided illumination mode. For protrusion monitoring experiments (Fig 4 A-C’’, Video 9) images were taken every 1 min for 10-14 hrs (Section 3.5.). Imaging started between 42 hpf and 48 hpf, except for LMNA overexpression which started at 48-53 hpf.

#### 2.2. Confocal scans

Fixed samples were imaged in a laser-scanning microscope (LSM 700 inverted, LSM 880 Airy upright; ZEISS) or point scanning microscope (2photon inverted; ZEISS) using the 40×/1.2 C-Apochromat water immersion objective (ZEISS). The samples were mounted in 1% agarose in glass-bottom dishes (MatTek Corporation) filled with E3 medium and imaged at room temperature. The microscopes were operated with the ZEN 2011 (black edition) software (ZEISS).

### 3. IMAGE PROCESSING AND ANALYSIS

#### 3.1. Sample drift correction

First, maximum projected sub-stacks (five *z* slices) of the raw live images were generated in Fiji. XY-drift of 2D stacks was then corrected using a manual drift correction Fiji plug-in created by Benoit Lombardot (Scientific Computing facility, Max Planck Institute of Molecular Cell Biology and Genetics, Dresden, Germany). The script can be found on (<u>imagej.net/Manual_drift_correction_plugin</u>). The drift corrected movies were then used for tracking migrating HCs.

#### 3.2. Tracking migrating HCs

The migrating HCs were manually tracked by following the centre of the cell body in 2D drift-corrected images using MTrackJ plug-in in Fiji (Meijering, Dzyubachyk and Smal 2012).

#### 3.3. Deconvolution

The raw LSFM data was deconvolved in ZEN 2014 software (black edition, release version 9.0) using the Nearest Neighbor algorithm. Minimal image pre-processing was implemented prior to image analysis, using open-source ImageJ/Fiji software (<u>fiji.sc</u>). Processing consisted of extracting image subsets or maximum intensity projections of a few slices. Processed files were analyzed in Fiji.

#### 3.4. Morphodynamic analysis of HC migration

To quantify HC cell and nuclear morphodynamics, the free and open-source platform for bioimage analysis Icy (de Chaumont et al. 2012, Manich et al. 202) (http://icy.bioimageanalysis.org) was used to automatically digitize cell and nuclear contours. The “Active Contours” plugin was used to segment the contours of HC cell and nuclear outlines during migration and in mitosis (Materials and Methods). Because the retina is densely-packed, the segmentation only worked when the cell of interest was singled out from the background and had enough distance from neighboring cells. To meet this goal, we used two different approaches: 1-cell transplantation (see section 1.7. Blastomere transplantations) and 2-DNA injection (see section 1.3.). A step-to-step manual of the protocols and plugins to measure cell and nucleus morphodynamics is available in (http://icy.bioimageanalysis.org) (Manich et al, 2020).

In this study, we extracted the following shape descriptors from Icy analysis: perimeter (µm), sphericity (%), elongation ratio (a.u.). “Perimeter” measures the perimeter of the region of interest (ROI) in micrometers. “Sphericity” is a measure of how similar to a sphere the ROI is. “Elongation ratio” is a scale factor given by the ratio between the first and second ellipse diameters of an ROI. The minimum value is 1 (for a round object).

#### 3.5. Protrusion tracking and analysis

Protrusions were analyzed using light-sheet time lapse video recording of *Tg(ptf1a:Gal4-VP16, UAS:gap-YFP)* at 1 min intervals. The protrusion tips were manually tracked simultaneously with HC centroids using the MTrackJ plug-in in Fiji (section 3.2.). The angle between the protrusion and the direction of HC movement was defined as the angle between the unit vector defined by the direction of HC movement and the unit vector pointing to the protrusion tip from the HC centroid. All plots for this part of the analysis were created in Matplotlib (Hutner, 2007).

#### 3.6. Utrophin fluorescence intensity distribution profiles

Utrophin fluorescence intensity distribution profiles of migrating HCs were measured in Fiji by drawing a line (width=3) along the cell axis at each time point. The utrophin signal intensity was measured using the max projection of 3 consecutive central z planes of the cell.

### 3. *In vivo* Brillouin light scatter microscopy

#### 3.1. Mounting of zebrafish larvae for in vivo Brillouin

Embryos were anesthetized in MS-222 (0.02% in E3; Sigma-Aldrich) for approximately 20 minutes and placed in a lateral position on a glass-bottom dish suitable for optical imaging. Some specimens were placed on a polyacrylamide gel that acted as a spacer between the glass bottom and the embryo (for gel preparation see (Schlussler et al. 2018). A drop (200 *μ*l) of low-gelling-point agarose (1% in E3, 30°C; Sigma-Aldrich) was used to immobilize the embryo. Immobilized larvae were then immersed in MS-222 (0.02%) and 1-phenyl-2-thiourea (PTU, 0.003%, Sigma-Aldrich) containing E3 during imaging. All embryos were released from the agarose embedding between Brillouin measurements and kept under standard conditions as described in Section 1.1.

#### 3.2. Brillouin microscopy set up and data analysis

The Brillouin shift measurements were performed using a custom-built Brillouin microscopy setup described in Schlüßler, Möllmert et al. 2018. The setup consists of a frequency-stabilized diode laser with a wavelength of 780.24 nm (DLC TA PRO 780; Toptica, Gräfelfing, Germany), a confocal unit employing a Plan Neofluar 20x objective (Carl Zeiss Microscopy; NA 0.5) and a two-stage virtually imaged phased array spectrometer. The setup was controlled using a custom program written in C++ (https://github.com/BrillouinMicroscopy/BrillouinAcquisition). The improvement of the Brillouin microscopy setup by adding a Fabry-Pérot interferometer (FPI) lead to the suppression of previously described strong reflections from the glass surface and rendered the gel spacer in subsequent measurements unnecessary. The data acquired was evaluated using a custom MATLAB (The MathWorks, Natick, MA) program (https://github.com/BrillouinMicroscopy/BrillouinEvaluation).

## Acknowledgments

We thank Otger Campàs, Rita Mateus, Teije Middelkoop, Carl Modes and Jaakko Lehtimäki for constructive feedback on project and manuscript. We are grateful to Heike Hollak and Sylvia Kaufmann for their technical support. R.A. thanks Stephan W. Grill for financial support and and Rita Mateus for generously providing lab space and reagents for experiments. We thank the Light Microscopy Facility of the Max Planck Institute of Molecular Cell Biology and Genetics (MPI-CBG), especially Jan Peychl, for their advice and technical support on image acquisition. We are also grateful to the Fish Facility of the MPI-CBG for technical assistance. We thank Gayathri Nadar from the Scientific Computing Facility at the MPI-CBG for discussions on data analysis. R.A. is grateful to Ivan Baines and Anthony Hyman for their help and support. R.S. thanks Simon Alberti for financial support.

R.A. was a research fellow of the Natural Sciences and Engineering Research Council of Canada (NSERC) from 2017-2019 and was a research fellow of the Fonds de la Recherche du Québec-Santé (FRQ-S) from 2019-2020. J.G. was supported by the Alexander-von-Humboldt Stiftung (Humboldt-Professorship), the European Commission through an ERC Starting Grant (‘‘LightTouch,’’ grant agreement number 282060) and the Deutsche Forschungsgemeinschaft (SPP 2191– Molecular mechanisms of functional phase separation, grant agreement number 419138906 to S.A. and J.G.). C.N. was supported by MPI-CBG, the FCG-IGC, Fundação para a Ciência e a Tecnologia Investigator grant (CEECIND/03268/2018) and an ERC consolidator grant (H2020 ERC-2018-CoG-81904).

## Author Contributions

R.A. and C.N conceived the study and designed experiments. R.A. performed all experiments and analyzed data. A.B. performed data analysis for protrusion angle distribution. R.S. realized the Brillouin microscopy, performed and evaluated the Brillouin measurements. R.A. and S.M. prepared the zebrafish specimens, acquired and evaluated the brightfield and confocal fluorescence images related to Brillouin microscopy. Data curation, making figures and movies for the manuscript was done by R.A. with input from C.N. R.A. and C.N. wrote the manuscript with input from all authors.

## FIGURE LEGENDS

**Supplementary Figure 1.**
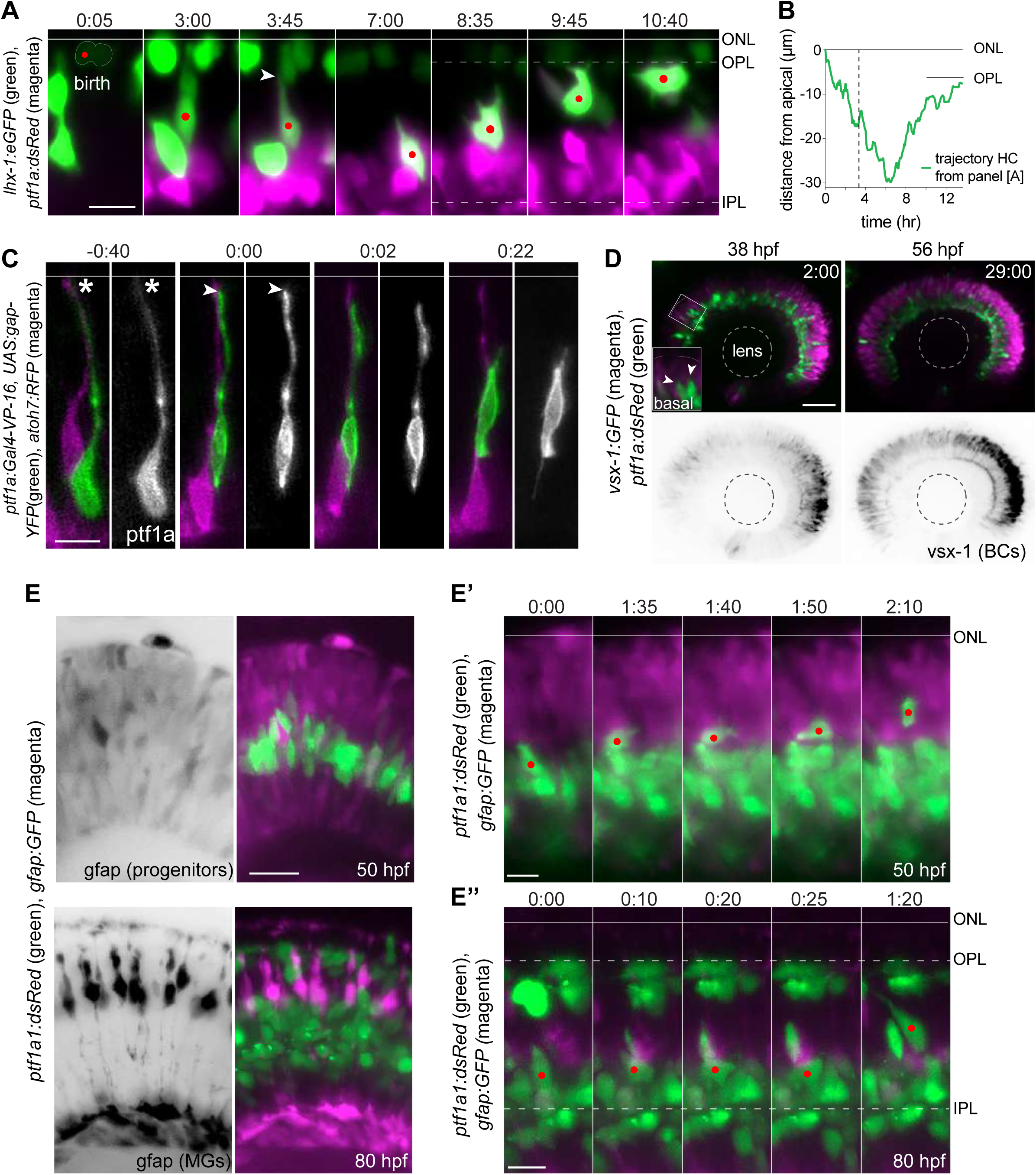
**(A)** Montage of bidirectional migration of an HC from birth to final positioning. Red dot: tracked HC; arrowhead: HC detachment from the apical surface. *Tg(lhx-1:eGFP)* labels HCs [green], *Tg(Ptf1a:dsRed*) marks ACs and HCs [magenta]. Scale bar: 10 µm. **(B)** Trajectory of migrating HC (depicted in A) from birth to terminal position relative to the apical surface; ONL (0 µm). **(C)** Retraction of HC apical attachment. *Tg(ptf1a:Gal4-VP-16, UAS:gap-YFP)* labels membrane of HCs [green], and *Tg(atoh7:RFP)* labels membranes of PRs and RGCs [magenta]. The labeled cell was tracked until it reached the HC layer. asterisk: apical attachment; arrowhead: tip of the retracted attachment; Line: apical surface. Scale bar: 10 µm. Note: HC was followed until final positioning. **(D)** Stills from time-lapse images of a retina before (38 hpf) and after (65 hpf) BC lamination. *Tg(ptf1a:dsRed)* labels HCs and ACs [green], *Tg(vsx1:GFP)* marks BCs [magenta]. Scale bar: 50 μm. Higher magnification inset of the outlined region shows two HCs moving apically [arrowheads]. **(E-E”)** HCs do not migrate along radially-oriented progenitors or MGs. *Tg(Ptf1a:dsRed*) marks ACs and HCs [green], *Tg(gfap:GFP)* labels MGs [magenta]. **(E) Top:** Prior to MG generation (50 hpf). GFAP^+^ cells are neurogenic progenitors. **Bottom:** Mature bipolar MG morphology (80 hpf). Scale bar: 20 µm. **(E’)** Montage of an HC migrating perpendicular to GFAP^+^ progenitors in 50 hpf, **(E’’)** stills of an HC migrating perpendicular to mature MGs in 80 hpf. Scale bar: 10 µm. Line: apical surface; red dot: migrating HC; dotted line: OPL (top), IPL (bottom). Time in h:min (A, C, E’-E’’).

**Supplementary Figure 2.**
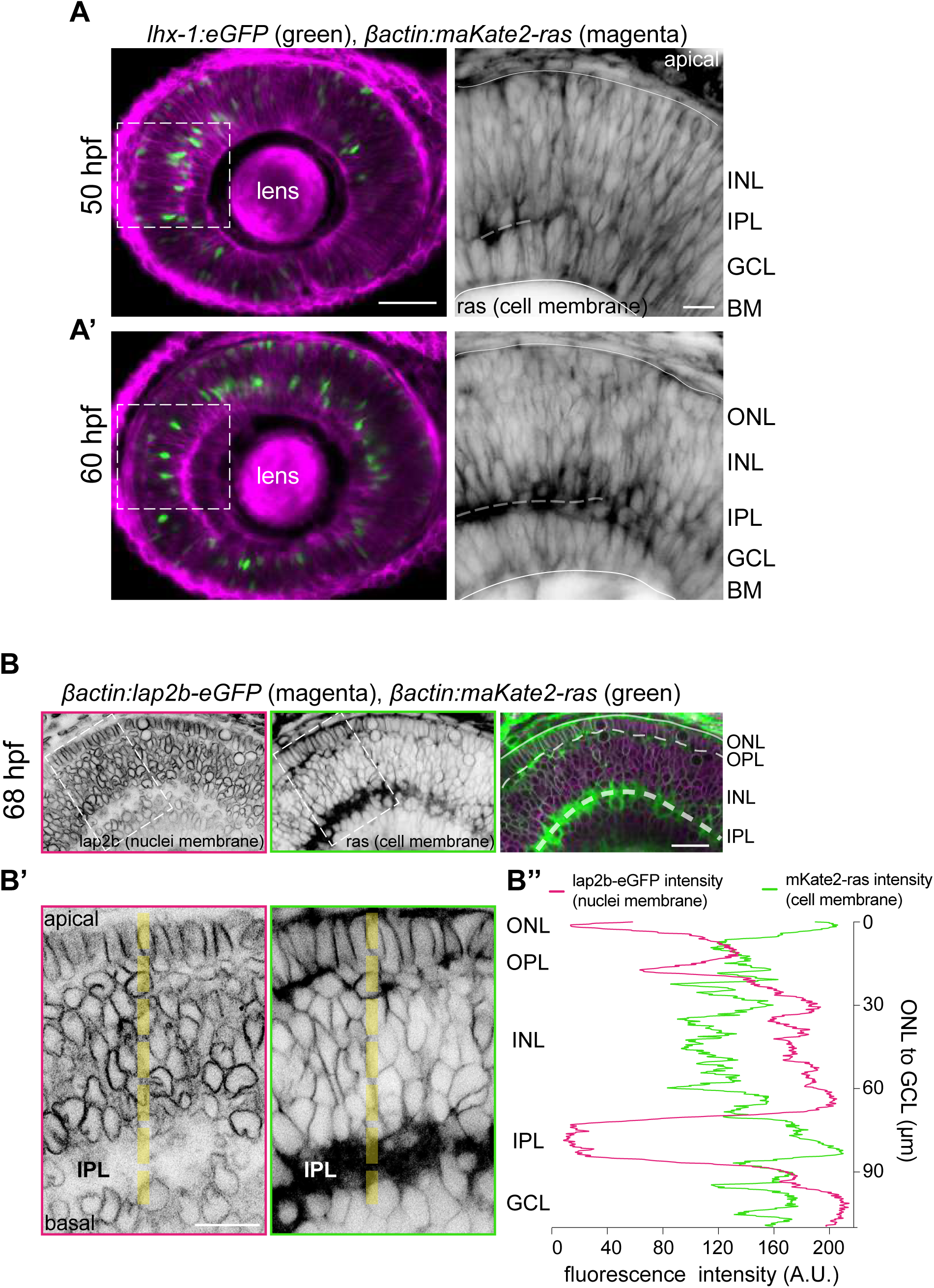
**(A)** Stills from time-lapse of a retina at 50 hpf (left panel) and 60 hpf (right panel). *Tg(lhx-1:eGFP)* labels HCs [green], *Tg(*βactin*:maKate2-ras)* marks cell membrane of all retinal cells [magenta]. Scale bar: 50 µm. Higher magnification inset of the outlined region show cell membranes. Arrowheads: forming IPL. Scale bar: 50 μm. **(B)** Confocal sections of *Tg(βactin:maKate2-ras)* labeling cell membrane of all retinal cells [green] and *Tg(βactin:lap2b-eGFP)* labeling nuclear envelope of all retinal cells [magenta], at 68 hpf. **(B’)** Higher magnification inset of the outlined region. Yellow dashed line: line scan measurements of lap2b-eGFP (nuclei membrane) and mKate2- ras (cell membrane), from ONL-to-GCL. **(B’’)** Average fluorescence intensity profiles of lap2b-eGFP (nuclei membrane) and mKate2-ras (cell membrane) from ONL-to-GCL. While, lap2b-eGFP (nuclei membrane) intensity profiles are the lowest at OPL and IPL, mKate2-ras (cell membrane) are highest. This shows that OPL and IPL are highly enriched with membrane. Scale bar: 20 µm.

**Supplementary Figure 3.**
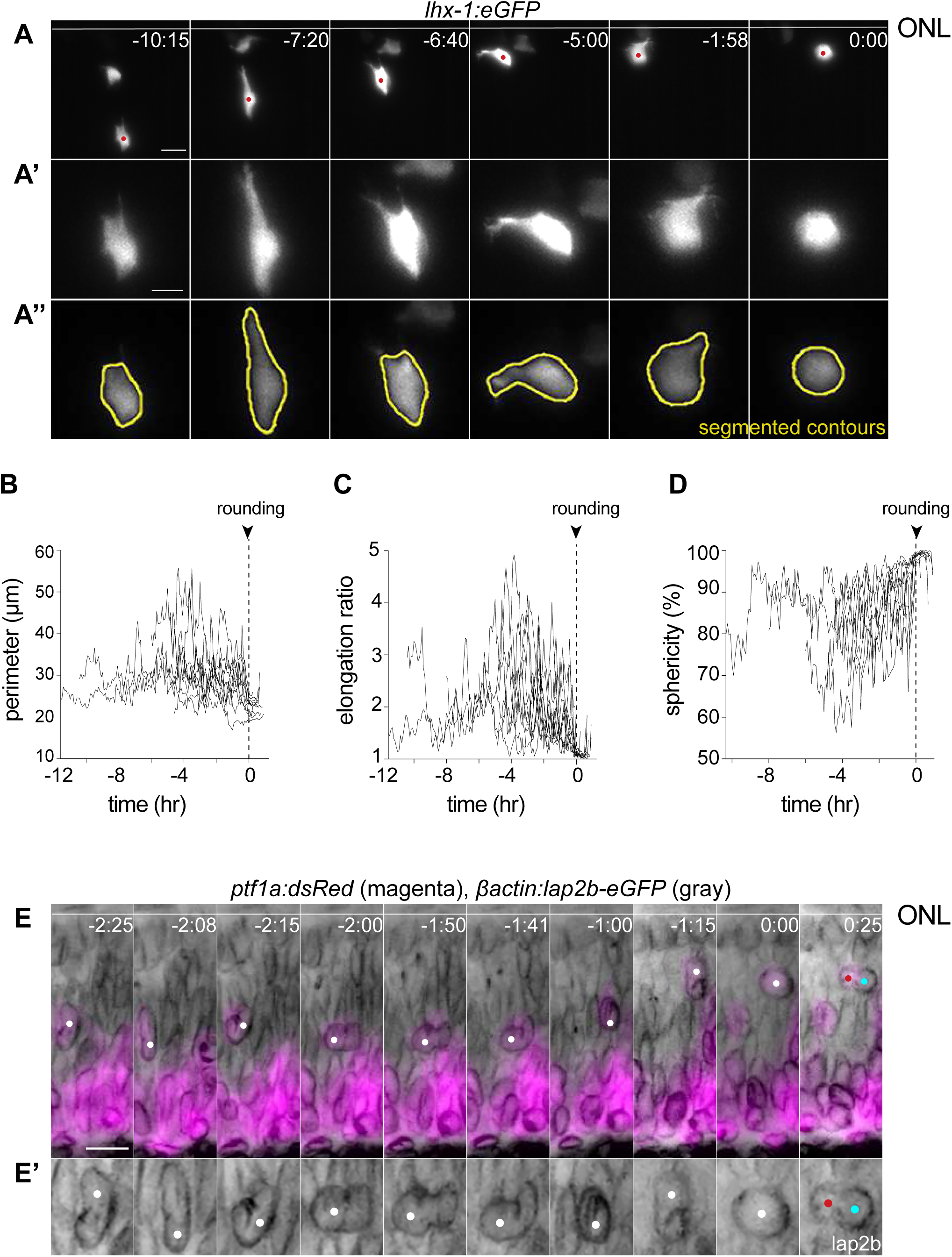
**(A)** Cytoplasmic deformations of an HC migrating from basal INL to the HC layer. Cells from *Tg(lhx-1:eGFP)* transgene embryo were transplanted into WT embryos. White line: ONL; red dot: migrating HC. Scale bar: 10 μm. **(A’)** Close-up of HC in (C). Scale bar: 5 μm. **(A’’)** Representative sequences of the automated segmented contours of the cell body (yellow line) generated by Icy. **(B-D)** Cell morphodynamic changes in 12 HCs from 7 embryos; **(B)** perimeter (μm), **(C)** elongation ratio, **(D)** sphericity (%). **(E)** HCs nuclei undergo nuclear deformations as they squeeze through the crowded retina. *Tg(*βactin*:lap2b-eGFP)* labels nuclear envelopes [gray]. *Tg(ptf1a:dsRed)* is expressed in HCs and ACs [magenta]. Line: ONL; white dot: migrating HC; red and cyan dots: HC sister cells after division. Scale bars: 10 μm. **(E’)** Higher magnification insets of HC nuclei envelope shown in the bottom panel. Scale bar: 5 μm. Time in h:min (A, E)

**Supplementary Figure 4.**
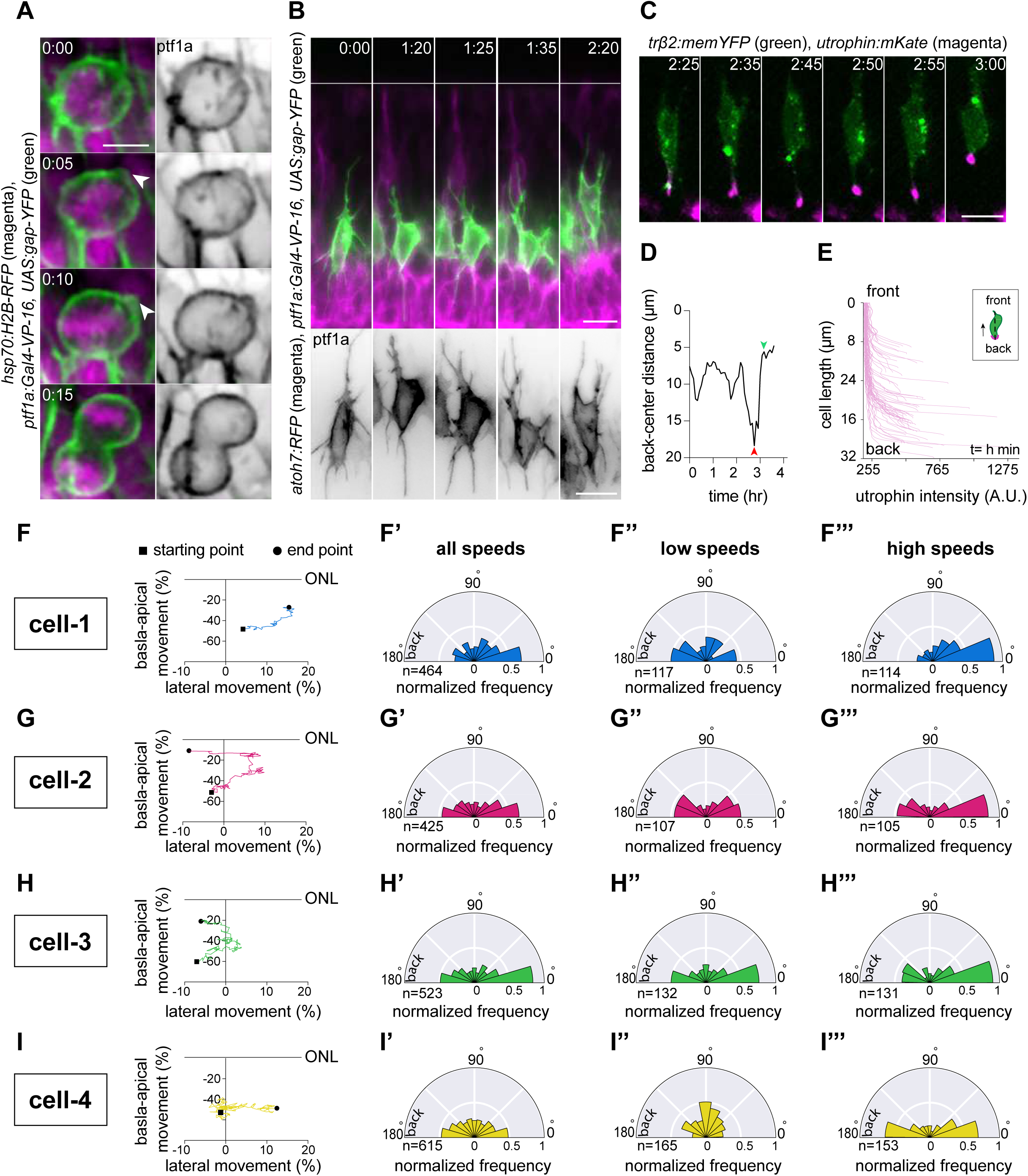
**(A)** Time-series of an HC during mitosis shows that HCs display membrane blebs (white arrowhead) during rounding. *Tg(ptf1a:Gal4-VP-16,UAS:gap-YFP)* labels HC membrane [green] and *Tg(hsp70:H2B-RFP)* is expressed in nuclei [magenta]. Scale bar: 5 μm. **(B)** Time-lapse sequence of dynamic protrusions in a migrating HC. Protrusions display different morphologies and point towards different directions. *Tg(ptf1a:Gal4-VP-16,UAS:gap-YFP)* labels HC membrane [green]. Scale bar: 20 μm. Bottom: close up of HC membrane [gray]. Scale bar: 10 μm. White line: ONL. **(C)** Time-series of a migrating HC [green] (labeled by trβ2:memYFP) and its dynamically changing uropod [magenta] (labeled by utrophin-mKate). **(D)** Measurements of the length of the uropod relative to HC centroid shows that it undergoes extension [red arrowhead] followed by retraction [green arrowhead]. **(E)** Average utrophin:mKate fluorescent intensity profile of images at (E). Time in h:min (A-F). **(F-I)** Migration trajectories and the frequency distribution of the angle between the direction of instantaneous HC movement and its protrusions in 4 HCs; at all speeds **(F’-I’)** at high speed: speeds above 75 percentile of the cell speed **(F’’-I’’)**, and at low speed: below 25 percentile of that cell’s speed **(F’’’-I’’’)**. A protrusion pointing exactly toward the direction of cell movement has an angle of 0°, and a protrusion pointing exactly opposite has an angle of 180°. The numbers represent number of angles observed in each category. The radius indicates the normalized frequency for each angle bin. i.e. the number of frames observed with the angle belonging to the particular angle bin for the given velocity condition normalized by the total number of frames observed in the given velocity condition.

**Supplementary figure 5.**
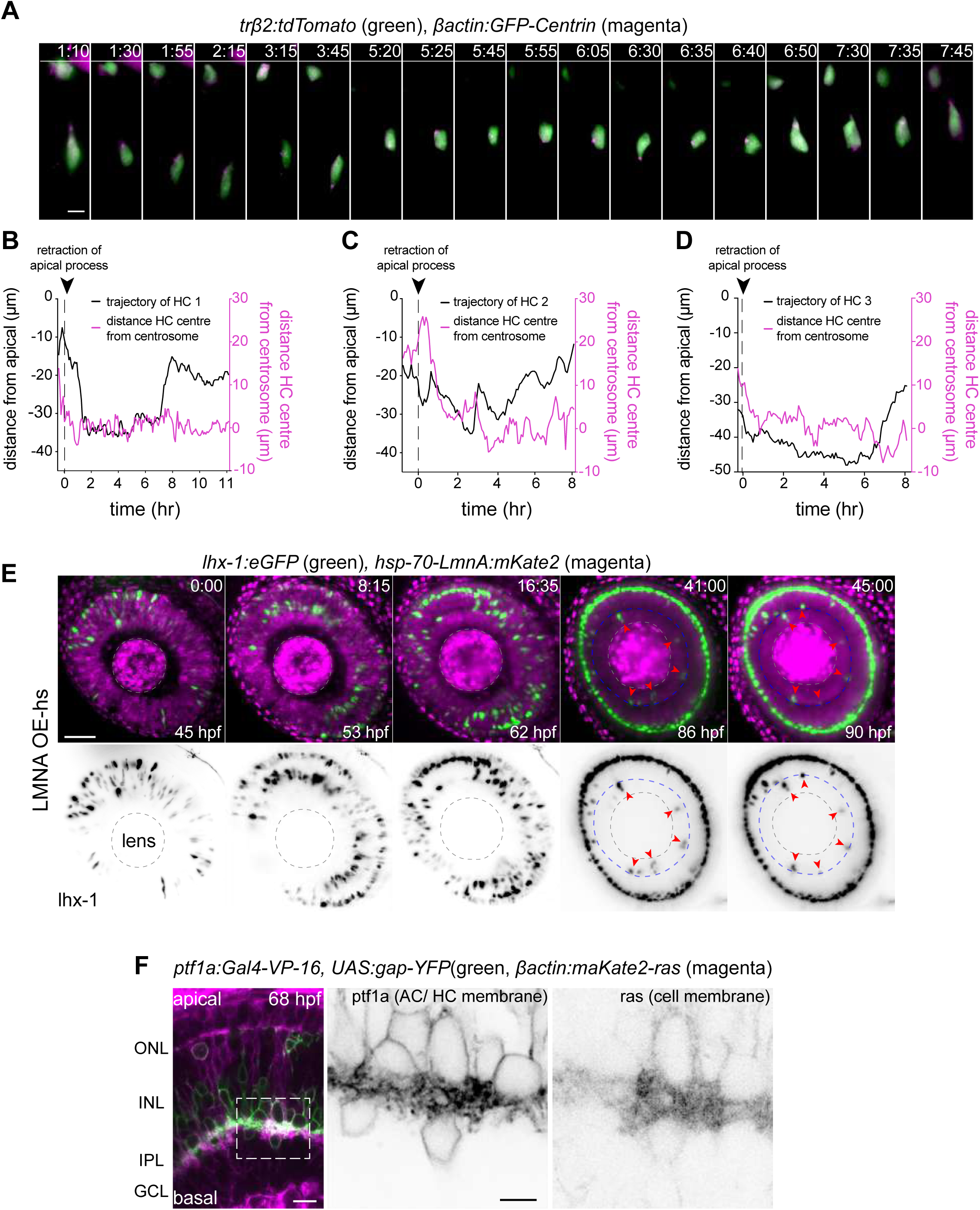
**(A)** Time-series showing the dynamics of centrosome position in a migrating HC. **(G)** bactin:GFP-Centrin and trb2:tdTomato DNA plasmids label centrosomes [magenta] and HCs [green], respectively. Scale bar: 20 μm. **(B-D)** Graphs showing migration trajectory of three migrating HCs (black line) and the distance between their centrosomes and center (magenta line) throughout migration. **(B)** The represented HC from (A). Arrowhead: time of detachment from the apical process; white line: ONL; dashed white line: HC layer. **(E)** Time-series of heat-shocked *Tg(lhx-1:eGFP) x Tg(hsp70:LMNA-mKate2)* double-transgene for 45 hr from 45 hpf shows that LMNA overexpression impairs proper HC layer formation. Red arrowheads: trapped HCs; dashed blue circle: IPL; dashed white circle: lens. Scale bar: 50 μm. **(E)** Confocal sections of *Tg(βactin:maKate2-ras)* labeling cell membrane of all retinal cells [magenta] and *Tg(ptf1a:Gal4-VP-16, UAS:gap-YFP)* labeling membrane and neurites of ACs and HCs [green], at 68 hpf. IPL is highly enriched in membrane and neurites. Higher magnification inset of the outlined region are on right. Scale bar: 10 μm.

## VIDEO CAPTIONS

**Video 1. Bidirectional trajectory of a tracked HC from apical detachment to final positioning.**

Time-lapse imaging of a typical bidirectional and bimodal migration of an HC. Upon retraction from the apical surface, HC enters a multipolar migration phase. Final mitosis occurs close to the HC layer. *Tg(lhx-1:eGFP)* labels HCs [green], *Tg(Ptf1a:dsRed*) marks ACs and HCs [magenta]. Red dot: tracked HC; white and blue dots: sister cells of the tracked HC after division. Time in h:min. Scale bar: 5 µm

**Video 2. Retraction of HC apical attachment.**

*Tg(ptf1a:Gal4-VP-16, UAS:gap-YFP)* labels membrane of HCs [green], and *Tg(atoh7:RFP)* labels membranes of PRs and RGCs [magenta]. White arrowhead: apical attachment; red arrowhead: tip of the retracted process. Time in h:min. Scale bar: 5 µm

**Video 3. Tangential migratory track of an HC after BC lamination.**

*Tg(vsx-1:GFP)* labels BCs [magenta] and *Tg(ptf1a:DsRed)* marks HCs [green]. White dot: tracked HC. Time in h:min. Scale bar: 20 μm

**Video 4. HCs undergo frequent and reversible cell morphodynamic changes during migration.**

Left: Time-lapse imaging shows basal-to-apical migration of an HC which was transplanted from *Tg(lhx-1:eGFP)* transgene embryo into WT embryos. HC rounds at the apical side (last frame).

Right: Representative sequences of the automated segmented contours of the cell body (yellow line) generated by Icy.

Red dot: tracked HC. Time in h:min. Scale bar: 10 μm

**Video 5. Nuclear deformations during HC migration**

Time-lapse imaging of an HC shows that it undergoes undergo nuclear deformations while squeezing through the crowded retina. βactin*:lap2b-eGFP* labels nuclear envelopes [gray], *trβ2:tdTomato* is expressed in HCs [magenta]. Black arrowhead: tracked HC. Time in h:min. Scale bars: 10 μm.

**Video 6. HC undergoes cell-shape deformations to bypass its neighboring cell.**

Time-lapse imaging of an HC migration from basal to apical INL. *Tg(Ptf1a:dsRed*) labels HCs and ACs. Red dot: tracked HC. Time in h:min. Scale bar: 10 µm.

**Video 7. HCs send dynamic protrusions during migration and only form blebs prior to mitosis.**

Time-lapse imaging of an HC migration shows that it sends dynamic protrusions while migrating. Upon rounding prior to mitosis, the tracked HC displays membrane-blebbing. *Tg(ptf1a:Gal4-VP-16,UAS:gap-YFP)* labels HC membrane. Red dot: tracked HC, red arrowhead: membrane-bleb, white and black dots: sister cells of the tracked HC after division. Time in h:min. Scale bar: 20 µm.

**Video 8. Angle distribution.**

The frequency distribution of the angle between the direction of instantaneous HC movement and its protrusions. The radius indicates the normalized frequency for each angle bin.

**Video 9. Lifeact-GFP is detected at the leading protrusions and the cell cortex of migrating HCs**

Left: trβ2:tdTomato marks HCs [green] and Lifeact-GFP labels all filamentous F-actin [magenta]. Time in h:min. Scale bar: 10 µm.

**Video 10. Migrating HCs display a front-rear utrophin distribution**

trβ2:memYFP is expressed in HCs [green] and utrophin-mKate marks the stable filamentous F-actin [magenta. Time in h:min. Scale bar: 10 µm.

## SUPPLEMENTARY TABLES

**S1 table:**
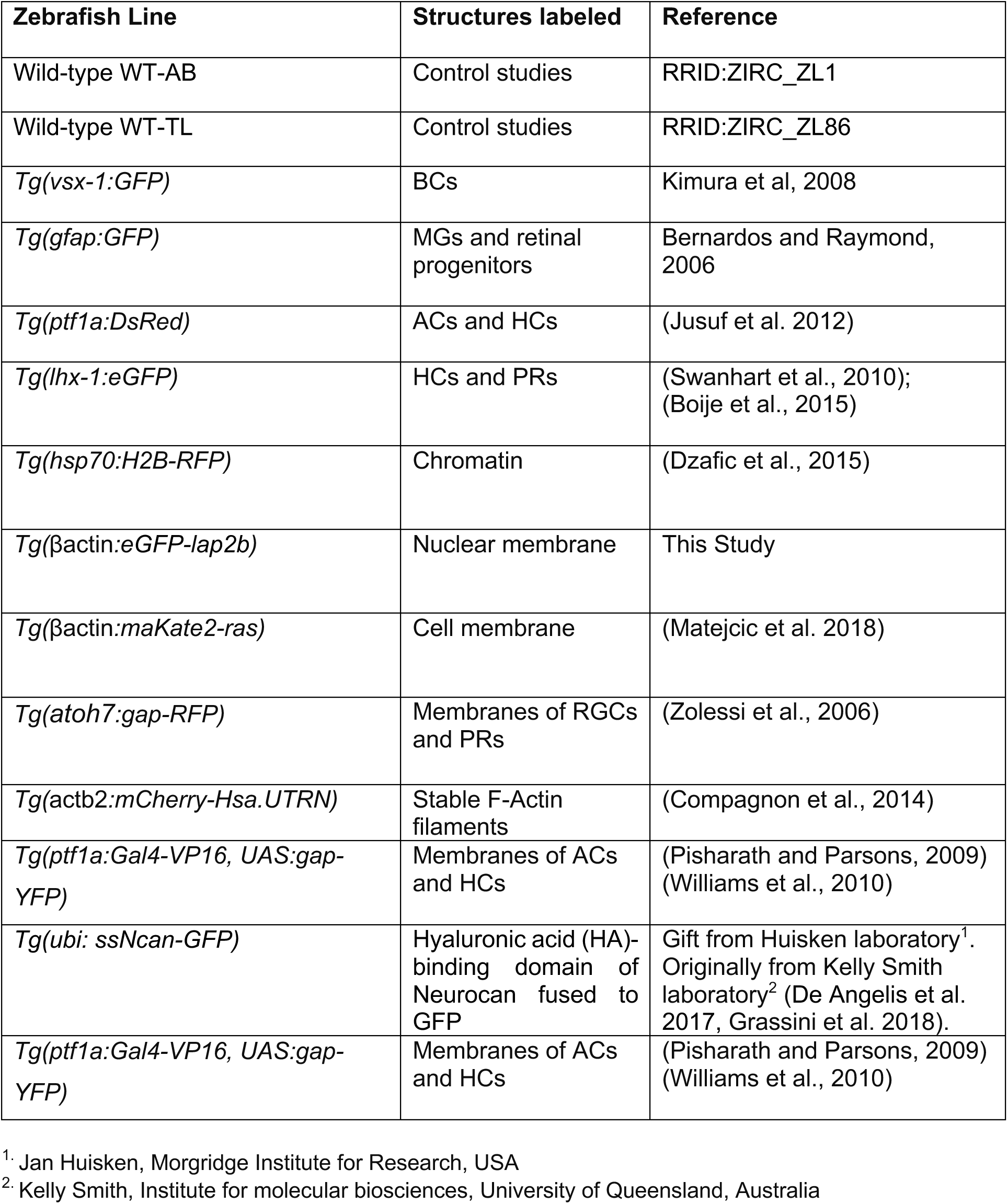
Zebrafish transgenic lines

**S2 table:**
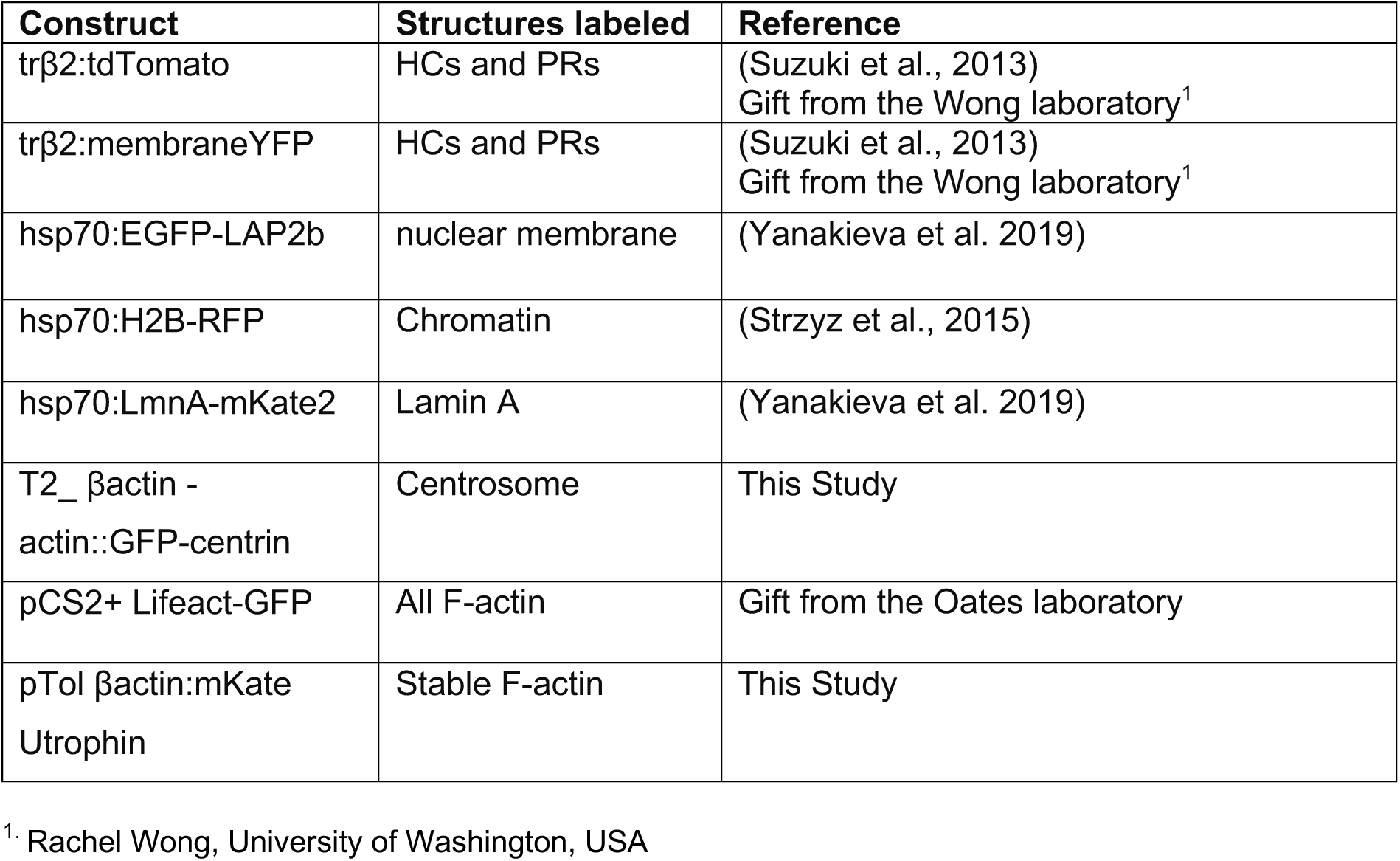
List of DNA constructs

**S3 table:** List of antibodies used for immunofluorescence

| Antibody | Structures labeled | dilution | RRIDs |
| --- | --- | --- | --- |
| Anti-Laminin | Laminin α1 | 1:100 | Sigma Aldrich, L9393,<br>RRID:AB_477163 |
| Zn-5 | RGCs | 1:50 | ZIRC, RRID:AB_10013770 |
| Alexa Fluor 647<br>Phalloidin | F-actin | 1:500 | Cell Signaling |
| DAPI | Chromatin | 1:1000 | Thermo Fisher Scientific |

**S4 table:** List of transgenes, DNA constructs, antibodies and their corresponding figures and videos

| **Zebrafish Line/ DNA construct/ antibody** | **Structures labeled** | **Figure/ Video** |
| --- | --- | --- |
| *Tg(vsx-1:GFP) x Tg(ptf1a:dsRed)* | BCs/ HCs, ACs | Fig 1C<br>Sup-Fig 1D<br>Video 3 |
| *Tg(gfap:GFP) x Tg(ptf1a:dsRed)* | MGs/ HCs, ACs | Fig 1E<br>Sup-Fig 1E-E'' |
| *Tg(lhx-1:eGFP) x Tg(ptf1a:dsRed)* | HCs, ACs and PRs | Fig 6A<br>Sup-Fig 1A<br>Video 1 |
| *Tg(ptf1a:Gal4-VP16, UAS:gap-YFP) x Tg(atoh7:gap-RFP)* | HCs, ACs/ RGCs, PRs | Sup-Fig 1C<br>Fig 4A-C''<br>Sup-Fig 4B<br>Video 2, Video 7 |
| *Tg(lhx-1:eGFP) + anti-Laminin* | HCs, ACs/ Laminin α1 | Fig 2A |
| *Tg(lhx-1:eGFP) x Tg(HA-GFP)* | HCs, ACs/ HA | Fig 2B |
| *Tg(lhx-1:eGFP) x Tg(atoh7:gap-RFP)* | HCs, PRs/ RGCs, PRs | Fig 2D-E''<br>Fig 6D |
| *Tg(lhx-1:eGFP) x Tg(βactin:maKate2-ras)* | HCs, PRs/ all membranes | Sup-Fig 2A-A' |
| *Tg(βactin:maKate2-ras) x Tg(βactin:lap2b-eGFP)* | All membrane/ all nuclei membranes | Sup-Fig 2B-B' |
| *hsp70:EGFP-LAP2b + trβ2:tdTomato plasmid* | nuclei membrane/ HCs, PRs | Fig 3A-A'' |
| *Tg(lhx-1:eGFP) x Tg(hsp70:H2B-RFP)* | HCs, PRs/ nuclei | Fig 3F-F' |
| *Tg(lhx-1:eGFP)*<br>Transplanted into wild-type | HCs, PRs | Sup-Fig 3A-A'' |
| *Tg(ptf1a:dsRed) + hsp70:EGFP-LAP2b plasmid* | HCs, ACs/ nuclei membrane | Sup-Fig 3E-E'<br>Video 5 |
| *pCS2+ Lifeact-GFP plasmid + trβ2:tdTomato plasmid* | All F-Actin filaments/ HCs, PRs | Fig 4D<br>Video 9 |
| *pTol βactin:mKate Utrophin plasmid + trβ2:membraneYFP plasmid* | Stable F-Actin filaments/ HCs, PRs | Fig 4E-F'<br>Sup-Fig 4C-E<br>Video 10 |
| *T2_ βactin::GFP-centrin plasmid + trβ2:tdTomato plasmid* | centrosome/ HCs, PRs | Fig 4D-E<br>Sup-Fig 4F-I |
| *Tg(ptf1a:Gal4-VP16, UAS:gap-YFP) x Tg(hsp70:H2B-RFP)* | Membranes of ACs and HCs/ nuclei | Sup-Fig 4A |
| *Tg(lhx-1:eGFP) + Phalloidin + DAPI* | HCs, PRs/ phalloidin/ DAPI | Fig 5A-B |
| *Tg(lhx-1:eGFP) x Tg(hsp70:LmnA-mKate2)* | HCs, PRs/ overexpressed LMNA | Fig 5B-D<br>Sup-Fig 5E |
| *Tg(lhx-1:eGFP) x Tg(ptf1a:DsRed) +*<br>Anti-Zn5 | HCs, PRs/ HCs, ACs/<br>RGCs | Fig 6A |
| *Tg(ptf1a:Gal4-VP16, UAS:gap-YFP) x*<br>*Tg(βactin:maKate2-ras)* | Membranes of ACs and<br>HCs/ membrane of all<br>retinal cells | Sup-Fig 5F |

